# Phototriggered profiling of receptor-proximal proteins in vivo in minutes

**DOI:** 10.1101/2023.05.25.542239

**Authors:** Mikiko Takato, Seiji Sakamoto, Hiroshi Nonaka, Tomonori Tamura, Itaru Hamachi

## Abstract

Neurotransmitter receptors are regulated by an extensive and dynamic network of protein-protein interactions. Understanding how these networks control synaptic transmission and give rise to higher-order brain functions necessitates their investigation in the live mammalian brain. However, tools available for this purpose lack the temporal resolution necessary to capture rapid changes in the interactome in live animals and require potentially disruptive genetic modifications to the animal under study. Here, we describe a method for light-activated proximity labelling in the live mouse brain that relies solely on small-molecule reagents and achieves a minute-order temporal resolution. Named PhoxID (photooxidation-driven proximity labelling for protein identification), this method involves selectively tethering a chemical photosensitiser to neurotransmitter receptors of interest and enabled us to identify characteristic as well as less studied interactors of the endogenous α-amino-3-hydroxy-5-methyl-4-isoxazolepropionic acid receptor (AMPAR) and the ψ-aminobutyric acid type A receptor (GABA_A_R) with just minutes of in-brain green light irradiation. Furthermore, PhoxID’s temporal precision allowed us to capture molecular snapshots of the AMPAR-proximal proteome in the postnatal developing cerebellum, leading to the discovery of age-dependent shifts. Overall, this work establishes a highly flexible and generalisable platform to study receptor interactomes and proximal microenvironments in genetically intact specimens with an unprecedented temporal resolution.

## Introduction

Neurotransmitter receptors play a central role in information processing in the central nervous system.^1^ Their activity and subcellular localisation, which dictate aspects of synaptic strength and plasticity,^2^ are heavily influenced by an elaborate and everchanging network of protein-protein interactions that is challenging to reconstitute in vitro. Thus, to decode neural circuits and higher-order brain functions at the molecular level, it is essential to study neurotransmitter receptor interactomes in the live, intact brain in its full complexity. Immunoprecipitation or affinity purification coupled with mass spectrometry (IP/AP-MS) has been widely used to identify proteins that complex with various neurotransmitter receptors.^3, 4^ However, it suffers from the loss of spatiotemporal information and the identification of spurious interactions that occur only in tissue homogenates. Moreover, it cannot capture weak or transient interactions that do not survive lysis conditions.^5^ In recent years, proximity labelling strategies have emerged as an alternative and valuable means to characterise protein interactomes in situ.^6–16^ Typically, engineered biotin ligases^8, 9^ or peroxidases^10, 11^ are expressed in cells as a fusion with a protein of interest. Upon addition of its substrate, the enzyme generates a highly reactive intermediate which can tag the proximal proteins. Peroxidases have been applied to map the excitatory and inhibitory synaptic cleft proteomes in cultured neurons^14^ but are challenging to deploy in vivo due to the toxicity of hydrogen peroxide. Biotin ligases, on the other hand, have been successfully applied to the live mouse brain, but they require biotin administration for days before the labelled proteomes can be harvested^15, 16^ and thus lack the temporal resolution necessary to distinguish proteomic changes that progress on shorter timescales. Moreover, enzyme-based proximity labelling methods in general require an exogenous enzyme to be fused to the protein of interest, which may perturb the structure, function, and trafficking of the native protein as well as obstruct biomolecular interactions that may otherwise occur.

To address these challenges, in recent years, we^17^ and others^18–25^ have developed chemistry-based, light-activated proximity labelling methods. These techniques use synthetic photocatalysts or photosensitisers tethered to affinity moieties for a target protein that can generate highly reactive species (carbenes^18, 19, 21^, phenoxy radicals^24^, singlet oxygen^17, 23, 25^) upon light irradiation. These nonenzymatic strategies allow proximity labelling to be carried out in genetically intact specimens, and the use of light as a trigger affords spatiotemporal control over the labelling reaction. However, the application of chemistry-based proximity labelling strategies to live animals is hampered by the difficulty of targeting synthetic probes to proteins of interest in vivo.

Here, we have overcome this challenge and established a workflow that enables the phototriggered profiling of neurotransmitter receptor proximal-proteomes in genetically intact live mice with just minutes of photoirradiation. Our method, named PhoxID (photooxidation-driven proximity labelling for protein identification), consists of two consecutive steps (**Fig. 1a**): In the first step, a small-molecule photosensitiser is covalently and selectively anchored to a target receptor in the live mouse brain by a designer synthetic molecule. In the second step, the photosensitiser is activated by in-brain visible light irradiation to locally generate singlet oxygen, which oxidises proteins within a theoretical radius of ∼70 nm^26^ around the photosensitiser. A nucleophilic probe is concomitantly administered to label the oxidised proteins with desthiobiotin, enabling their enrichment and identification by mass spectrometry. Following our workflow, we were able to profile the extracellular protein neighbourhoods of two representative neurotransmitter receptors, AMPARs and GABA_A_Rs, in vivo. Utilising PhoxID’s high temporal resolution, we were also able to capture molecular snapshots of the AMPAR-proximal proteome as it evolved during postnatal cerebellar development.

**Fig. 1.**
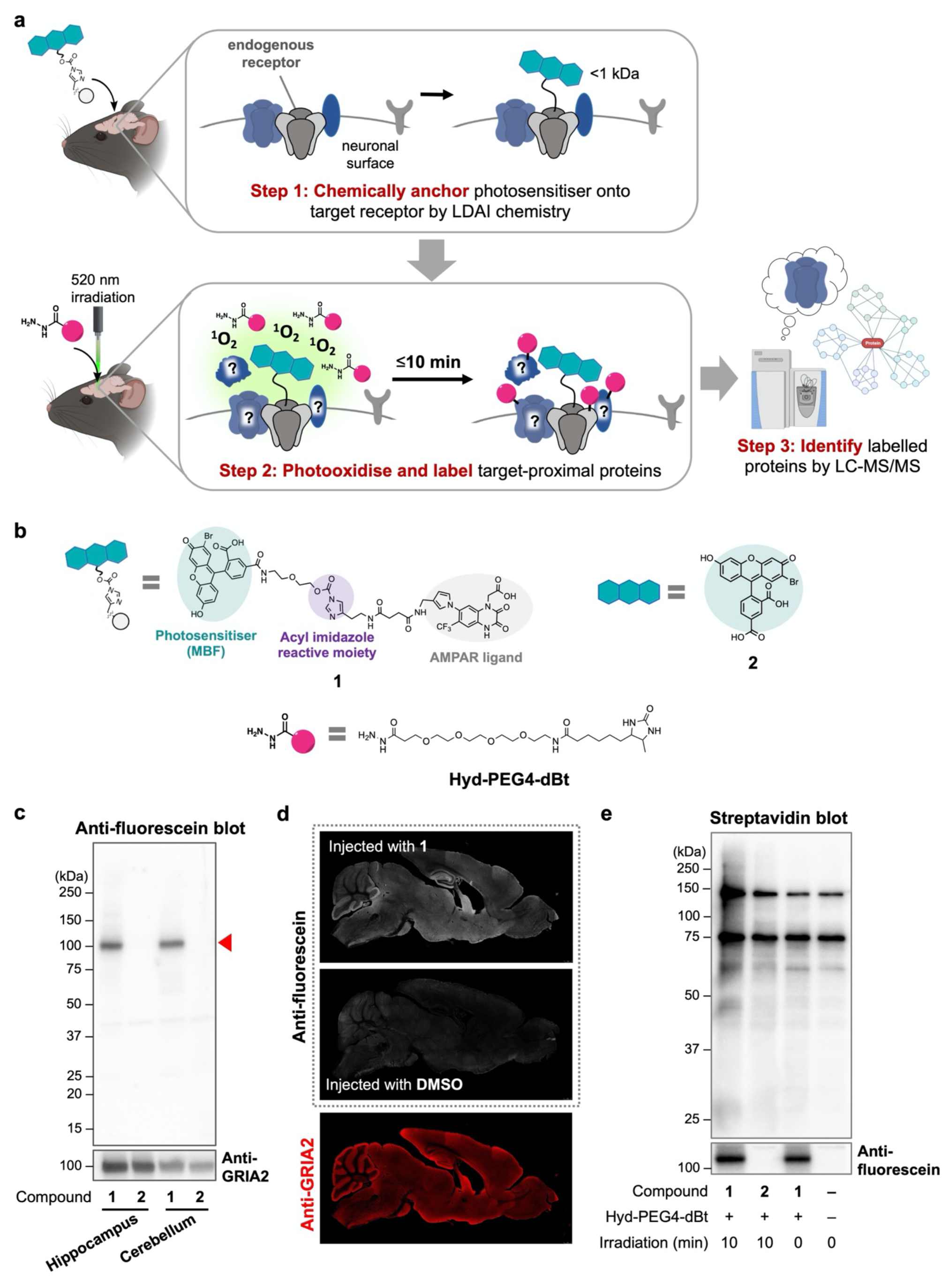
PhoxID enables light-triggered proximity labelling in the live mouse brain. **(a)** Illustration of the mechanism of PhoxID in the live mouse brain**. (b)** Chemical structures of the AMPAR-targeted photosensitiser **1,** the untargeted negative control photosensitiser **2,** and the labelling reagent Hyd-PEG4-dBt**. (c)** Western blot of brain tissue isolated from mice injected to the lateral ventricles with **1** and probed with anti-fluorescein, which recognises MBF. Red triangle indicates the band corresponding to AMPAR subunits covalently modified with MBF. **(d)** Confocal laser scanning microscopy images of brain slices prepared from mice injected to the lateral ventricles with **1** and stained with anti-fluorescein or anti-GRIA2. Scale bar, 1 mm. **(e)** Streptavidin blot of cerebellum proteins isolated and enriched from mice subjected to PhoxID.

## Results

### Establishing PhoxID in vivo for the identification of AMPAR-proximal proteins

For our initial trials, we aimed to label and identify the protein neighbours of AMPARs. Formed as heteromeric tetramers of four subunits (GRIA1−GRIA4), AMPARs are ionotropic glutamate receptors that play an essential role in synaptic plasticity and memory formation in the brain.^27^ However, the details of their transient extracellular interaction network remain elusive, as current knowledge about the AMPAR interactome is limited to IP/AP-MS studies of brain homogenates.^3, 28^ To chemically anchor a photosensitiser onto AMPARs in the live mouse brain for PhoxID, we employed ligand-directed acyl imidazole (LDAI) chemistry, a covalent protein labelling strategy developed by us.^29, 30^ Our recent study showed that this technique can be implemented in live mice to selectively incorporate desired probes into the extracellular ligand-binding domain of endogenous AMPARs.^31^ We thus leveraged LDAI chemistry to covalently tether synthetic photosensitisers to AMPARs in vivo. To this end, we synthesised the AMPAR-targeted LDAI reagent **1** (**Fig. 1b**) according to the design principle established in our previous studies.^29^ For the photosensitiser, we selected 2-monobromofluorescein (MBF), which we found generates singlet oxygen with a moderate quantum yield (Φ = 0.17 in EtOH) (**Extended Data Fig. 1**) upon excitation with green light (α_ex_ = 506 nm in aqueous solution). **2** was synthesised as an untargeted negative control reagent (**Fig. 1b**). To assess its specificity towards AMPARs, we injected **1** (100 µM, 4.4 µL) into the lateral ventricles of a C57BL/6N mouse (5 weeks old) and isolated the hippocampus and cerebellum tissues 24 h after injection. A unique advantage of the covalent anchoring method was that the selectivity of MBF attachment to AMPARs could be confirmed by western blot analysis of the tissue homogenates (**Fig. 1c**). Imaging by confocal microscopy also showed discernible MBF signal in the hippocampus and the molecular layer of cerebellum, where AMPARs are highly expressed (**Fig. 1d**).^32^ We also considered employing 4,5-dibromofluorescein (DBF) as the photosensitiser, since it possesses a higher singlet oxygen quantum yield (Φ = 0.29 in EtOH) ^33^ than MBF and has proven to be effective for photoactivated proximity labelling.^17^ However, we found that the anchoring of DBF onto AMPARs in vivo was considerably less efficient than that of MBF (**Extended Data Fig. 2**), presumably because the hydrophobic nature of the additional bromine atom resulted in low tissue dispersibility of the reagent. Thus, MBF was chosen as the photosensitiser for subsequent PhoxID experiments.

**Fig. 2.**
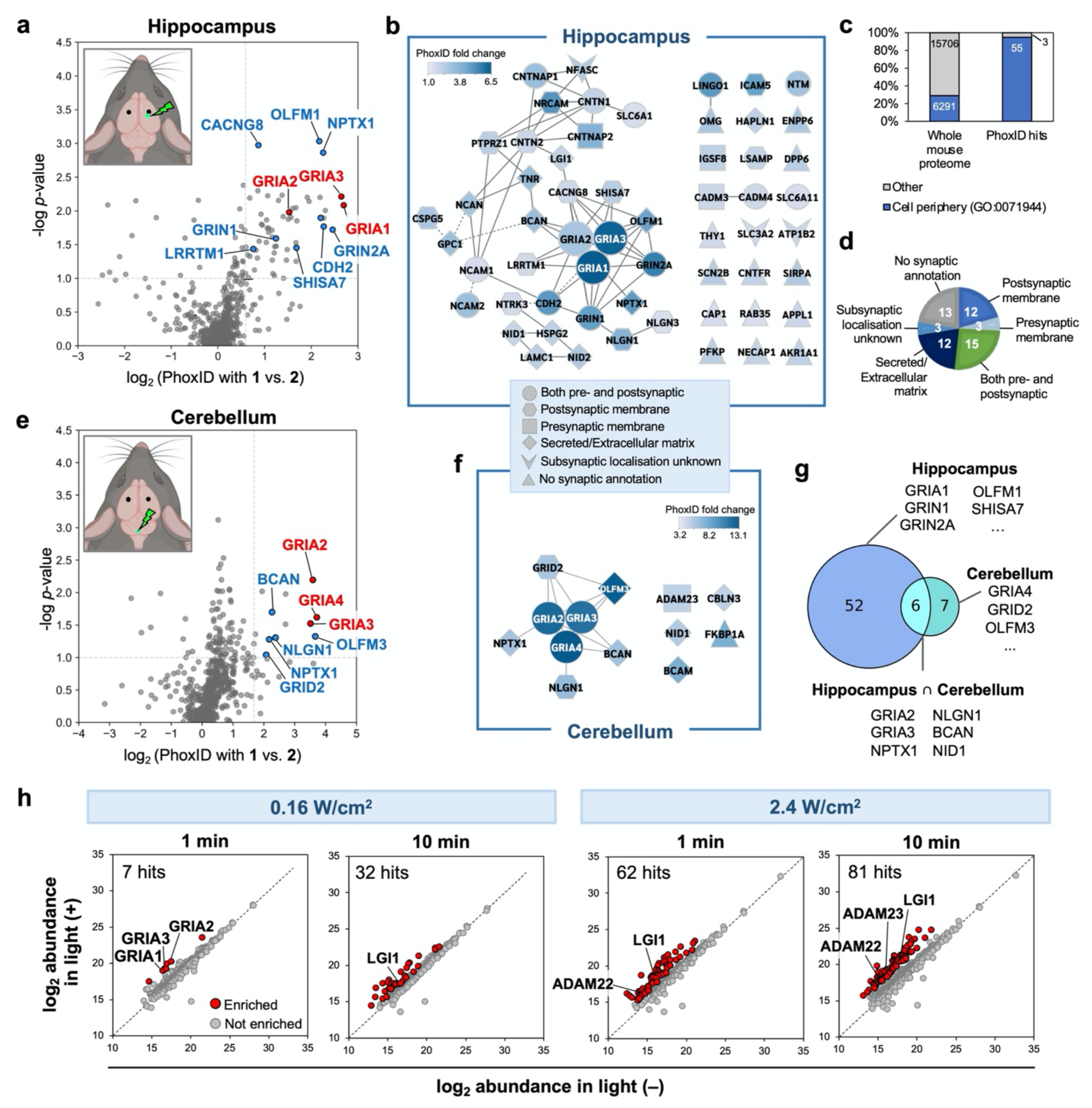
PhoxID can profile the AMPAR-proximal proteome in the live mouse brain in minutes. **(a, e)** Volcano plots of proteins identified by AMPAR-targeted PhoxID in (a) the hippocampus and (e) cerebellum (*n*=3 biological replicates). Fold changes are calculated from the extent of labelling in mice injected with **1** vs. **2**. Red and blue points represent AMPAR subunits and well-known AMPAR-interacting proteins, respectively. Dashed lines indicate the thresholds set for the data. Upper-left panels, schematic diagram of the brain injection sites. Black dots represent the injection sites of **1** or **2** and lime green dots represent the site of **Hyd-PEG4-dBt** injection and photoirradiation. **(b, f)** Network diagrams of known physical (solid edges) and functional (dashed edges) interactions between the hit proteins in (b) the hippocampus and (f) cerebellum. Interaction information was retrieved from STRING 11.5 and supplemented with manual annotations (see Supplementary Tables 1 and 2). Node shapes correspond to the reported subsynaptic localisation in the Gene Ontology Cellular Component (GOCC) database supplemented with several manual annotations (see Supplementary Table 1). Node colours are scaled to PhoxID fold change. **(c)** Bar graph showing the fraction of hit proteins that are annotated to the cell periphery relative to the whole mouse proteome recorded in the PANTHER knowledgebase. **(d)** Pie chart of the subsynaptic distribution of hit proteins. **(e)** Volcano plot of proteins identified by AMPAR-targeted PhoxID in the cerebellum. **(g)** Venn diagram showing the overlap between the hit proteomes of the hippocampus and cerebellum. **(h)** Scatter plot showing the degree of labelling in the hippocampus of mice subjected to photoirradiation of varying intensity and duration.

For the protein labelling reagent, we selected hydrazide-PEG4-desthiobiotin (**Hyd-PEG4-dBt**) (**Fig. 1b**) for its high nucleophilicity and cell-impermeability, ^23, 34^ which allow us to exclusively label the extracellular neighbourhoods of AMPARs. To confirm if photoinduced protein labelling could proceed in the brain using **Hyd-PEG4-dBt**, a mouse administered with **1** was injected 24–28 h later with it at the cerebellum (5 mM, 4.4 µL) and subsequently subjected to irradiation at the same site for 10 min with light (520 nm, 0.16 W/cm^2^) delivered through an optical fibre. Streptavidin blotting of the cerebellum tissue confirmed that promiscuous protein labelling occurred in the live mouse brain in a MBF-anchoring and photoirradiation-dependent manner (**Fig. 1e**). Minimal background labelling was observed in the absence of **1**, suggesting that light at 520 nm does not activate endogenous photosensitisers (such as flavins and porphyrins) to a significant extent (**Fig. 1e**).

Encouraged by these results, we proceeded to identify the labelled proteins by mass spectrometry. For this purpose, we injected a mouse with **1** or **2** into the lateral ventricles and carried out photoinduced labelling in the right hippocampus the following day (**Fig. 2a**). After irradiation, the hippocampus was quickly isolated and the labelled proteins were enriched, digested, and identified by nanoLC-MS/MS. Based on our threshold criteria (**Supplementary Note**), 58 proteins were classified as “hit” proteins that were significantly more labelled in mice treated with **1** than in mice treated with **2** (**Fig. 2a, b, Supplementary Table 1**). Of the hits, 95% were cell-peripheral proteins (**Fig. 2c**), confirming that labelling was largely restricted to the cell surface and extracellular space. No correlation was observed between the fold change of the hit proteins and their reported total abundance in the brain (**Extended Data Fig. 3a**), and proteins across a wide range of abundances were enriched (**Extended Data Fig. 3b**). Among the most highly enriched proteins were subunits of AMPAR itself, with GRIA1, GRIA2, and GRIA3, but not GRIA4, being enriched. This is consistent with reports that GRIA4 comprises only a small fraction of AMPAR subunits in the hippocampus (**Fig. 2a**).^3, 32^ Protein network analysis showed that multiple proteins known to directly interact with AMPARs at hippocampal glutamatergic synapses were among the hits, namely OLFM1, NPTX1, CACNG8, CDH2, LRRTM1, and GRIN1/2A. Pre- and postsynaptic proteins that localise to glutamatergic synapses but have not been reported to interact directly with AMPARs (**Fig. 2b, d**) were also enriched, and proteins with a known synaptic annotation accounted for nearly 60% of the hit proteins. Secreted proteins and extracellular matrix proteins made up another ∼20% (**Fig. 2d**), among which was brevican (BCAN), a proteoglycan that is believed to be involved in AMPAR trafficking from extrasynaptic regions to synapses.^35^ Together, these results confirmed that PhoxID can capture AMPAR-proximal proteins in the live mouse brain with spatial specificity in just minutes of photoirradiation.

**Fig. 3.**
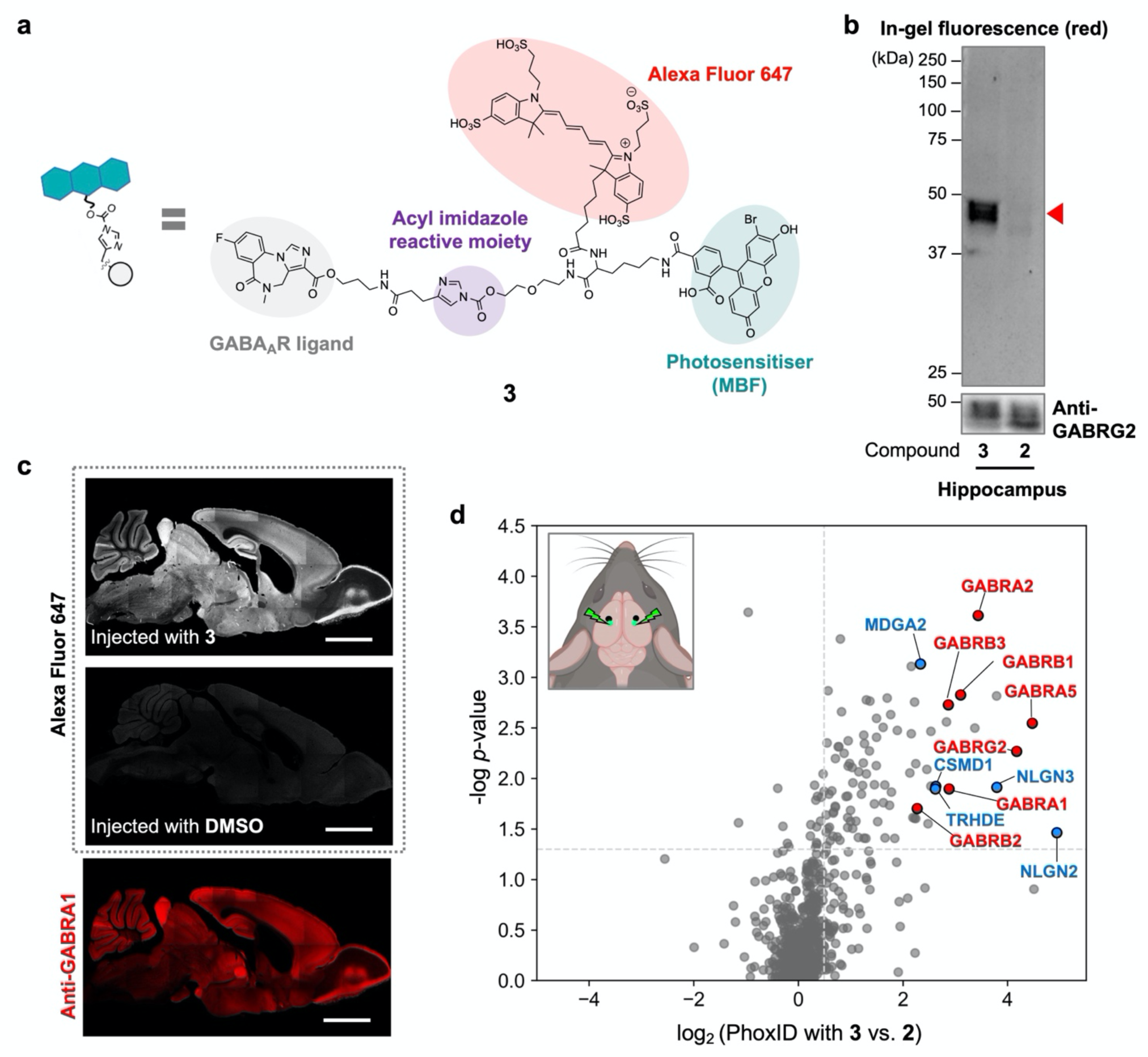
Expansion of PhoxID to the GABA_A_R-proximal proteome. **(a)** Chemical structure of the GABA_A_R-targeted photosensitiser **3. (b)** In-gel fluorescence analysis of hippocampus tissue from mice injected to the lateral ventricles with **3** or **2**. Red triangle points to the band corresponding to MBF/Alexa Fluor 647-modified GABA_A_Rs. **(c)** Confocal laser scanning microscopy imaging of Alexa Fluor 647 fluorescence observed in whole brain slices prepared from mice injected to the lateral ventricles with **3** or DMSO. Scale bar, 2 mm. **(d)** Volcano plot of GABA_A_R-targeted PhoxID in the hippocampus. Fold changes are calculated from the extent of labelling in mice injected with **3** vs. **2** (*n*=3 biological replicates). Red points represent GABA_A_R subunits. Blue points indicate GABAergic synapse proteins-of-interest. Dashed lines indicate the thresholds set for the data. Upper-left panel, schematic diagram of the brain injection and photoirradiation sites. Black dots represent the injection sites of **3** or **2** and lime green dots represent the site of **Hyd-PEG4-dBt** injection and photoirradiation.

We next asked whether PhoxID could shed light on regional differences in the AMPAR interactome. By simply changing the site of **Hyd-PEG4-dBt** administration and photoirradiation to the cerebellum, we were able to identify 13 proteins as hits there (**Fig. 2e, f, Supplementary Table 2**). In contrast to the hippocampus dataset, GRIA4 was identified in the cerebellum, where it is reported to account for over 60% of AMPAR subunits.^3^ We were also able to identify cerebellum-specific AMPAR interaction partners such as the secreted protein OLFM3 and the ionotropic glutamate receptor GRID2, which regulates AMPAR trafficking at parallel fibre (PF)-Purkinje cell synapses (**Fig. 2f, g**).^36^ These results demonstrate the utility of PhoxID in identifying components of the AMPAR-proximal proteome that vary by brain region.

We also investigated whether the extent of labelling in vivo can be tuned by adjusting the intensity and/or duration of photoirradiation (**Fig. 2h, Supplementary Table 3**). With 1 min of irradiation of the hippocampus at 0.16 W/cm^2^, only 7 proteins were identified as hits, 3 of which were subunits of AMPAR itself. However, when the irradiation intensity was raised 15-fold to 2.4 W/cm^2^, 62 proteins, including more AMPAR interactors and glutamatergic synapse proteins, could be identified in the same 1 min timeframe (**Extended Data Fig. 4a**), showing that the extent of PhoxID labelling can be controlled without compromising the temporal resolution. Further extension of light exposure to 10 min (2.4 W/cm^2^) resulted in even broader coverage of the AMPAR vicinity. For example, all components of the transsynaptic ADAM22-LGI1-ADAM23 complex that regulates AMPAR-mediated synaptic transmission could be identified under this condition,^37^ whereas it was only partially detected at lower light doses (**Fig. 2h, Supplementary Table 3**). We also found that the fold change value of the individual hit proteins increased with light duration and intensity (**Extended Data Fig. 4b**). In total, these data clearly show the PhoxID labelling range can be modulated by altering the light irradiation conditions.

**Fig. 4.**
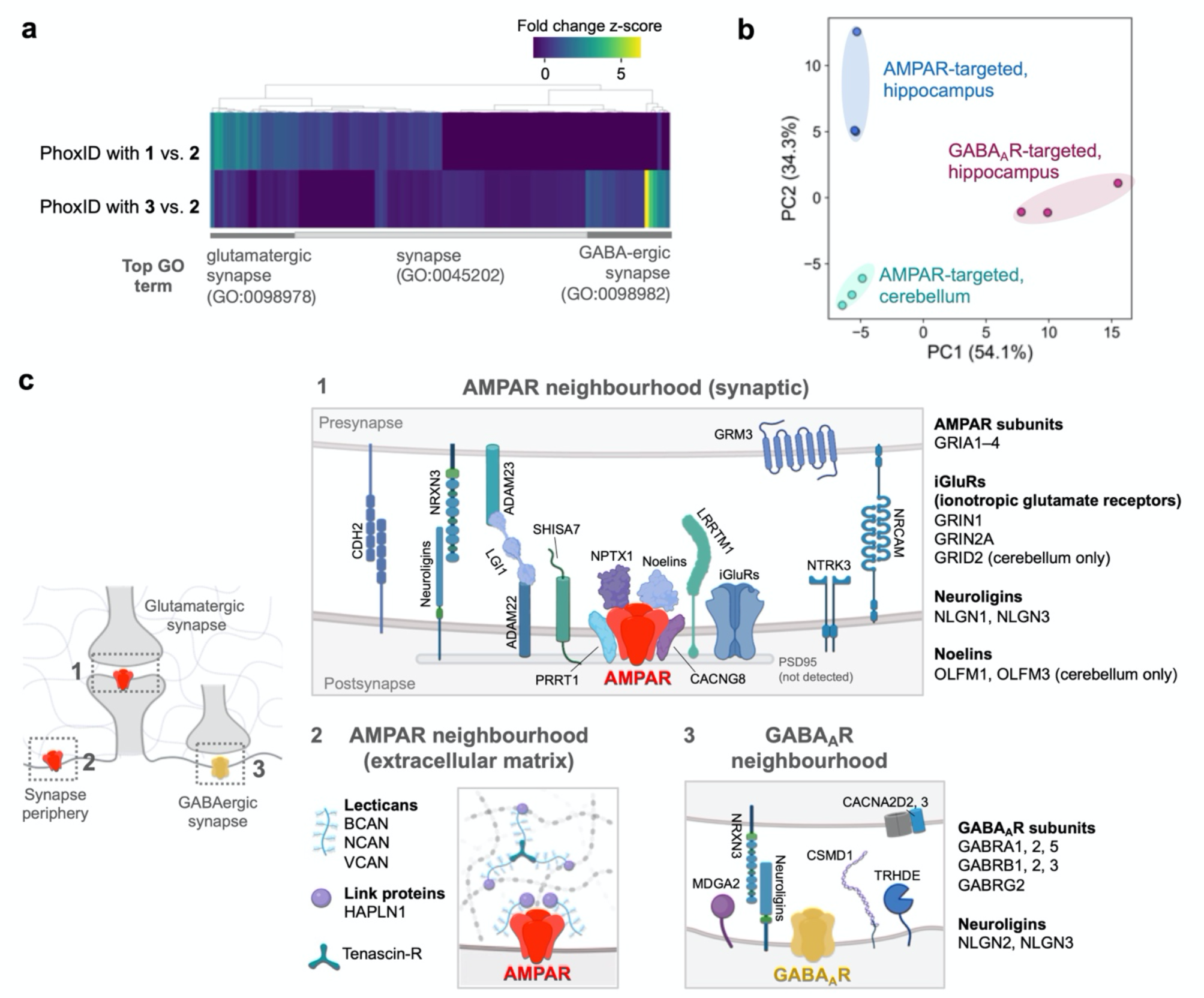
PhoxID can distinguish different neurotransmitter receptor neighbourhoods. **(a)** Hierarchically clustered heatmap of hit proteins identified in PhoxID with **1** and **3**. The top GOCC term based on raw *p*-value for the leftmost and rightmost 20 proteins, as well as all other proteins, are listed below the heatmap. Mitochondrial/nuclear contaminant proteins are omitted. **(b)** Principal component analysis of the AMPAR-proximal proteomes in the hippocampus and cerebellum, and the GABA_A_R-proximal proteome in the hippocampus, as identified by PhoxID. **(c)** Cartoon depiction of select proteins identified by in vivo PhoxID under the conditions described in Fig. 2 and 3. Panels 1 and 2, some AMPAR-proximal proteins known to conventionally be found in glutamatergic synapses and the extracellular matrix, respectively. Panel 3, some GABAergic synapse proteins identified by PhoxID.

### Application of PhoxID to the GABA_A_R-proximal proteome

We next asked whether PhoxID could characterise the protein neighbourhoods of other neurotransmitter receptors. We targeted the GABA_A_R, which is the chief mediator of inhibitory neurotransmission and whose protein microenvironment we expected to vary greatly from that of AMPARs.^38^ To attach MBF to GABA_A_Rs, we synthesised the LDAI reagent **3** comprised of the GABA_A_R ligand flumazenil connected to MBF via a linker branched with the hydrophilic fluorophore Alexa Fluor 647 to compensate for the hydrophobicity of flumazenil (**Fig. 3a**). ^31^ The design of **3** is such that Alexa Fluor 647 and MBF would be bound to GABA_A_Rs together, allowing us to infer the presence of MBF from Alexa Fluor 647 fluorescence. We injected **3** into the lateral ventricles of the mouse brain and verified that it could selectively and covalently anchor MBF onto GABA_A_Rs in the mouse hippocampus (**Fig. 3b**). Imaging by confocal microscopy also showed that Alexa Fluor 647 fluorescence overlapped with the localisation of GABA_A_Rs in the brain (**Fig. 3c**).^39^

After confirming the successful anchoring of MBF to GABA_A_Rs, we carried out PhoxID in the left and right hippocampi (**Fig. 3d, Supplementary Table 4**). Of the top 20 proteins with the highest fold changes (p<0.05), 7 were of subunits of GABA_A_R itself and at least 7 others had a known annotation to GABAergic synapses. The most highly enriched protein was NLGN2, a synaptic organiser molecule that is present exclusively at inhibitory synapses.^40^ Together, the enrichment of these proteins validated the labelling of the GABA_A_R precinct. Several proteins with scarce evidence supporting their localisation at inhibitory synapses were also in the hit protein list, namely CSMD1, TRHDE, and MDGA2 (**Fig. 3d**). These proteins were annotated to the inhibitory synapse for the first time by peroxidase-based proximity labelling in neuronal culture.^14^ Our results support the proposition that these proteins occupy inhibitory synapses, and do so in vivo.

We next compared the PhoxID datasets obtained for the AMPAR- and GABA_A_R-proximal proteomes. Hierarchical clustering analysis revealed that the hippocampal proteomes labelled by **1** and **3** diverged greatly, with **1** overwhelmingly labelling glutamatergic synapse proteins and **3** mainly enriching GABAergic synapse proteins (**Fig. 4a**). Principal component analysis (PCA) of the hippocampal AMPAR, cerebellar AMPAR, and hippocampal GABA_A_R hit proteomes clearly showed that PhoxID can distinguish the protein microenvironments of different target proteins and brain regions (**Fig. 4b**). **Fig. 4c** provides an illustration of notable AMPAR-proximal and GABA_A_R-proximal proteins identified as hits in this study, showing their localisation and diverse topologies. It should be noted that SHISA7 was detected only in AMPAR-targeted PhoxID. Although controversy has arisen in recent years as to whether SHISA7 interacts with AMPARs or GABA_A_Rs,^41, 42^ our proteomic data suggest that this protein is present in the vicinity of AMPARs in the mouse hippocampus.

Overall, these data show that PhoxID can distinguish and characterise proteomic landscapes spatially focused on neurotransmitter receptors of interest in vivo with high specificity.

### PhoxID can capture snapshots of the AMPAR-proximal proteome during postnatal development

During postnatal development, the neuronal circuitry in the cerebellum is structurally and functionally refined by eliminating redundant synapses while retaining and strengthening others.^43, 44^ We thought that PhoxID, with its minute-order temporal resolution, would be able to capture “snapshots” of the AMPAR-proximal protein territory in vivo during neonatal growth and reveal how it evolves during this critical time. Specifically, we decided to capture PhoxID snapshots of mice at postnatal day 8 (P8) (when PF synapse formation and climbing fibre synapse elimination have just begun), day 13 (P13) (when PF synapses are about to shift toward elimination), and week 5 (early adulthood) (**Fig. 5a**).^44^

**Fig. 5.**
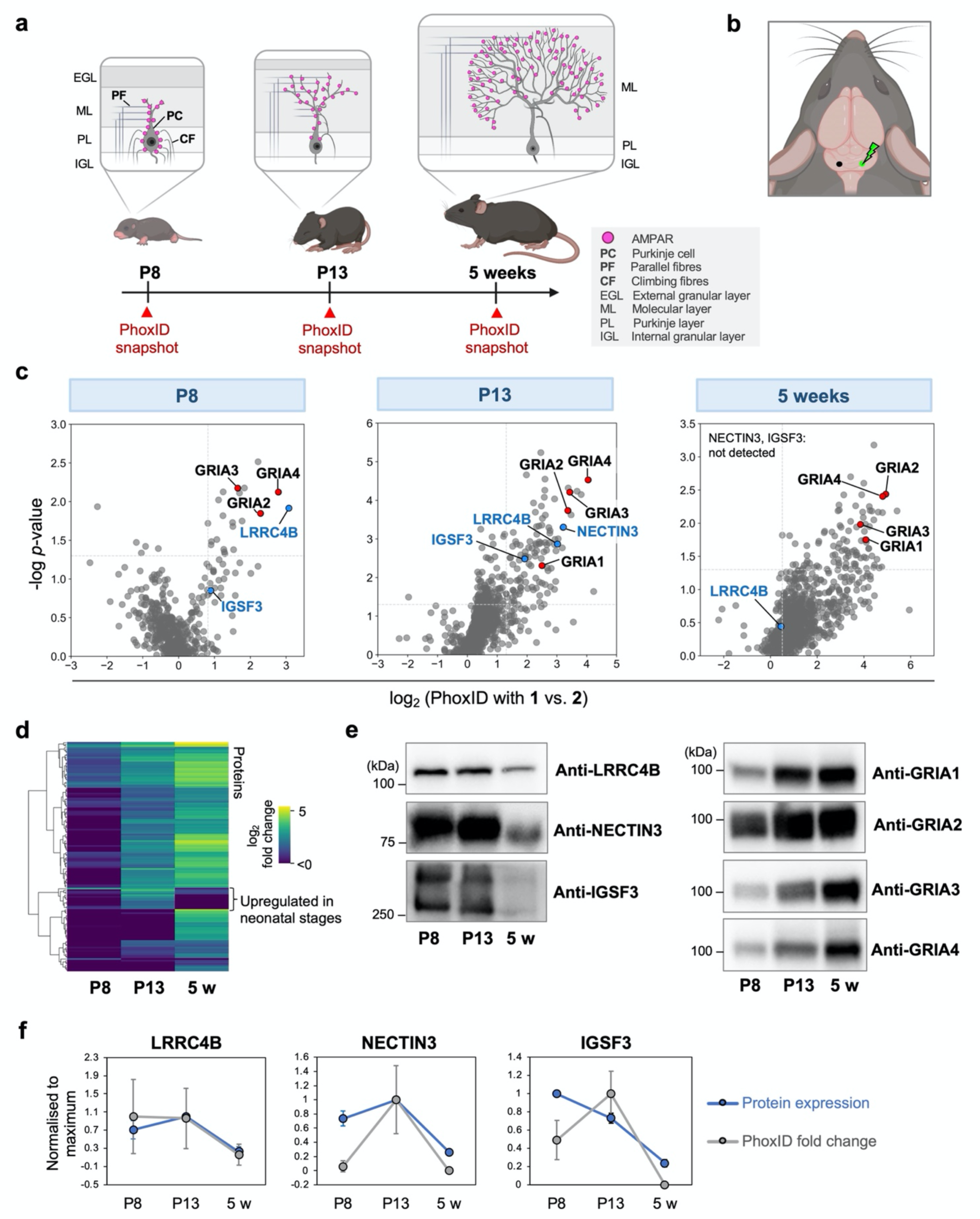
PhoxID reveals shifts in the AMPAR-proximal proteome in postnatal development. **(a)** Cartoon depicting the localisation of AMPARs in the mouse cerebellum during postnatal development. In vivo PhoxID was carried out on mice at P8, P13, and 5 weeks (5 w). **(b) 1** and **Hyd-PEG4-dBt** were injected to the left and right halves of the cerebellum, respectively. **(c)** Volcano plots of the PhoxID experiments conducted at each neonatal age studied (*n*=3 biological replicates). Proteins whose unique peptide count was <2 are not shown. NECTIN3 is not shown at P8 because only 1 unique peptide was detected. **(d)** Hierarchically clustered heatmap of PhoxID hits at P8, P13, and 5 weeks revealed a subset of proteins whose extent of labelling was higher at neonatal ages than at 5 weeks. Fold change values are shown for proteins that were classified as hits in at least one of the ages. Mitochondrial/nuclear contaminant proteins are omitted. **(e)** Western blot analysis of the expression levels of LRRC4B, NECTIN3, IGSF3, and the four AMPAR subunits at P8, P13, and 5 weeks. **(f)** Comparison of the PhoxID fold change of LRRC4B, NECTIN3, and IGSF3 at P8, P13, and 5 weeks, and their protein expression levels as determined in **(e)**. Data are normalised to the maximum value in each dataset. PhoxID fold changes are shown as median ± s.d. (*n*=3 biological replicates). Protein expression levels are shown as mean ± s.d. (*n*=3 technical replicates).

Juvenile mice have smaller brain volumes and significantly fewer AMPARs than at maturity, and their lateral ventricles are also underdeveloped. Thus, we sought to maximise labelling by directly injecting **1** into the cerebellum and carrying out **Hyd-PEG4-dBt** injection and photoirradiation (10 min, 0.16 W/cm^2^) at a proximal site (**Fig. 5b**). Following this protocol, AMPAR-proximal cerebellum proteomes were successfully obtained from mice at all ages (**Fig. 5c, Extended Data Fig. 5a, 6, Supplementary Table 5**). Relatively few proteins were enriched at P8, possibly because (1) AMPARs are poorly expressed in the early postnatal period and/or (2) there are fewer proteins near AMPARs due to immature spine formation. The PhoxID fold changes of most proteins generally increased with age (**Fig. 5d**), which likely was in part due to the abundance of MBF-modified AMPARs increasing with growth (**Extended Data Fig. 5b, c**). However, there were 14 proteins that opposed this trend and were more enriched in the AMPAR precinct at P8/P13 than at 5 weeks, suggestive of their age-dependent proximity to AMPARs (**Fig. 5d**) (see **Supplementary Table 5** for a full list). Of these, we focused on LRRC4B, NECTIN3, and IGSF3 for follow-up analysis because of their high detection confidence. All three of these proteins have been linked to synaptogenesis in the brain, ^45–48^ but their spatial relationship with AMPARs in the context of postnatal development is not known, warranting further study.

We first asked whether the cerebellum expression levels of these proteins were developmentally regulated. Western blot analysis revealed that while AMPAR expression steadily increased from P8 to 5 weeks, LRRC4B expression decreased approximately 2-fold during the same period (**Fig. 5e, f, Extended Data Fig. 5d, e**). PhoxID fold change levels of LRRC4B followed a similar trend, with enrichment levels being 6-fold higher in neonatal stages than in adulthood. Similar to LRRC4B, NECTIN3 expression also dropped dramatically from P8/P13 to week 5. Despite its substantial expression at P8, however, the PhoxID fold change of NECTIN3 at that age was very small. This discrepancy between the protein expression level and PhoxID fold change was also observed for IGSF3: The PhoxID fold change of IGSF3 increased from P8 to P13, while protein expression decreased monotonically from P8 to week 5.

Given that the PhoxID fold change of the hit proteins are expected to roughly reflect a combination of their average physical distance from AMPARs and their average abundance in the singlet oxygen diffusion range, we hypothesised that AMPARs reside in more LRRC4B-rich environments at P8 and P13 than at week 5, and in more NECTIN3- and IGSF3-rich environments at P13 than at P8 and week 5. To test this hypothesis, we attempted to confirm the localisation and distribution of these proteins in the cerebellum by immunostaining. Although the available antibodies for LRRC4B and IGSF3 failed to detect their targets under our conditions, a NECTIN3 antibody could specifically visualise endogenous NECTIN3 in the cerebellum (**Extended Data Fig. 7**). This imaging analysis showed that NECTIN3 is expressed throughout the cerebellum at P8 and P13, including in the molecular layer where AMPARs are expressed. By week 5, however, NECTIN3 signal in the molecular layer significantly diminished, confirming that AMPARs inhabit more NECTIN3-rich environments at P13 than at week 5. We are unsure as to why NECTIN3 was not enriched in P8 PhoxID despite its high expression in the molecular layer. One possibility is that PhoxID labelling was less effective in P8 mice compared to P13 mice due to the lower expression of AMPARs. An alternative theory is that the PhoxID results may be reflective of a change in localisation on the AMPAR side: At P8, we observed a substantial pool of AMPARs on Purkinje cell bodies, presumably at climbing fibre synapses^44^, in a relatively NECTIN3-poor environment. By P13, however, this pool is largely translocated to the dendritic arbors weaving through the NECTIN3-rich molecular layer.

In the adult brain, NECTIN3 is primarily found at puncta adherens junctions on hippocampal dendritic branches and serves as a “latch” to stabilise the synapses to which they are adjacent.^46, 47^ Interestingly, in the neonatal hippocampus, NECTIN3 has been reported to transiently dwell in synaptic junctions as they mature into dendritic spines.^46^ The localisability of NECTIN3 at synapses as suggested by these reports, combined with our PhoxID results, support the hypothesis that AMPARs populate NECTIN3-rich environments at P13.

Although we were unable to verify their localisation in the cerebellum by immunostaining, there is evidence in the literature supporting LRRC4B’s and IGSF3’s proximity to AMPARs. LRRC4B is an excitatory synapse protein known to promote bidirectional synapse differentiation through its interaction with the presynaptic leukocyte antigen-related family proteins and its docking to the postsynaptic scaffolding protein PSD-95, which also anchors AMPARs.^45^ Thus, LRRC4B and AMPARs are expected to colocalise. ^45^ Our PhoxID results provide the first clues that cerebellar LRRC4B expression, in the context of its proximity to AMPARs and in general, may be developmentally regulated. As for IGSF3, it is a little-studied protein that has been reported to transiently express on developing granule cell membranes, notably at developing PF-Purkinje cell contacts (where IGSF3 and AMPARs are mainly present on the PF and Purkinje cell sides, respectively). ^48^ Thus, it is possible that IGSF3 and AMPARs approach each other for a brief period of time before these junctions mature into synapses.

Collectively, these results illustrate how PhoxID’s high temporal resolution enables the hypothesis-free identification of ephemeral yet dynamic shifts in neurotransmitter receptor-proximal proteomes in vivo during neural development.

## Discussion

Studying neurotransmitter receptors and their interactomes in the genetically unedited, live brain, where neuronal circuits are intact and readily linked to behavioural outputs, is essential to deciphering high-order brain functions and pathologies at the molecular level. We have developed an entirely chemical, light-driven proximity labelling technology that is compatible with live animal studies for this purpose. At present, only genetically encoded tools have been successful at proximity labelling in vivo, and they suffer from a low temporal resolution on the order of hours to days. As we have shown herein, our method relies solely on small-molecule reagents and required as brief as 1 min to sufficiently label and characterise receptor-proximal proteomes, an unprecedented temporal resolution for an in vivo proximity labelling method. This enabled us to identify transitory shifts in the proteome that may be obscured by methods that can only obtain cumulative information over several days. The use of light as a trigger also made PhoxID easily customisable in terms of tissue region (adjustable by changing the optical fibre insertion site) and effective labelling radius (adjustable by tuning photoirradiation duration and intensity, making target ID to broader proteomics possible). In addition, the photosensitiser used in this study is excited by green light, which minimised background labelling caused by endogenous photosensitisers, a major problem for light-based proximity labelling approaches requiring blue excitation.^49^

It is important to note that PhoxID may overrepresent proteins with multiple histidine residues, the primary locus of singlet oxygen-mediated hydrazide labelling.^23^ Also, because we aimed to exclusively label the extracellular milieu of neurotransmitter receptors, this study was biased by design towards detecting proteins with large extracellular domains (**Extended Data Fig. 8**). To identify the intracellular interactome as well as increase the coverage of transmembrane proteins, a cell-permeable labelling reagent could be implemented in the future.^17^ In addition, the present study used a covalent method to anchor MBF to neurotransmitter receptors, but a noncovalent binding strategy would also be feasible with a specific nanobody or small-molecule ligand provided its dissociation rate is sufficiently low. Furthermore, although not discussed in detail here, we have also shown that PhoxID is compatible with studies in acute brain slices (**Extended Data Fig. 9, Supplementary Table 6**), demonstrating the robustness of PhoxID across specimen types. Finally, in the future, PhoxID could potentially be coupled with optogenetics or disease models to map proteomic changes associated with brain activity or pathological processes. In summary, we expect that PhoxID can be adapted to survey the interactome of various protein targets in a wide range of biological contexts, and serve as a powerful tool for exploring local proteomes as they rapidly change in place and time.

## Acknowledgements

The authors would like to thank K. Uchida, M. Ishikawa, Y. Yabuki and E. Kusaka for their support in chemical synthesis and compound characterisation; K. Nishimura and K. Matsuba for mass analysis; K. Nishizawa and T. Gonda for mouse experiments; H. Utsunomiya for early-stage evaluation of photosensitisers. This work was supported by the Japan Science and Technology Agency (JST) ERATO Grant JPMJER1802, a Grant-in-Aid for Specially Promoted Research (23H05405), and the JST CREST Grant JPMJCR1854 to I.H.; a Grant-in-Aid for Scientific Research on Innovative Areas ‘Integrated Bio-metal Science’ (19H05764), Scientific Research (B) (21H02058) to T.T.; and a Grant-in-Aid for JSPS Research Fellow (21J23228) to M.T.; the Science and Technology Platform Program for Advanced Biological Medicine (AMED/MEXT, grant number JP22am0401006) to H.N. The MS raw data and analysis files have been deposited in the ProteomeXchange Consortium (http://proteomecentral.proteomexchange.org) via the jPOST partner repository (http://jpostdb.org)50 with data set identifier JPST002162.

## Author Contributions

M.T., T.T., and I.H. conceived the project and designed the experiments. M.T., S.S., H.N., and T.T. performed the experiments and data analysis. M.T., T.T., and I.H. wrote the manuscript with input from all authors.

---

**Extended Data Fig. 1.**
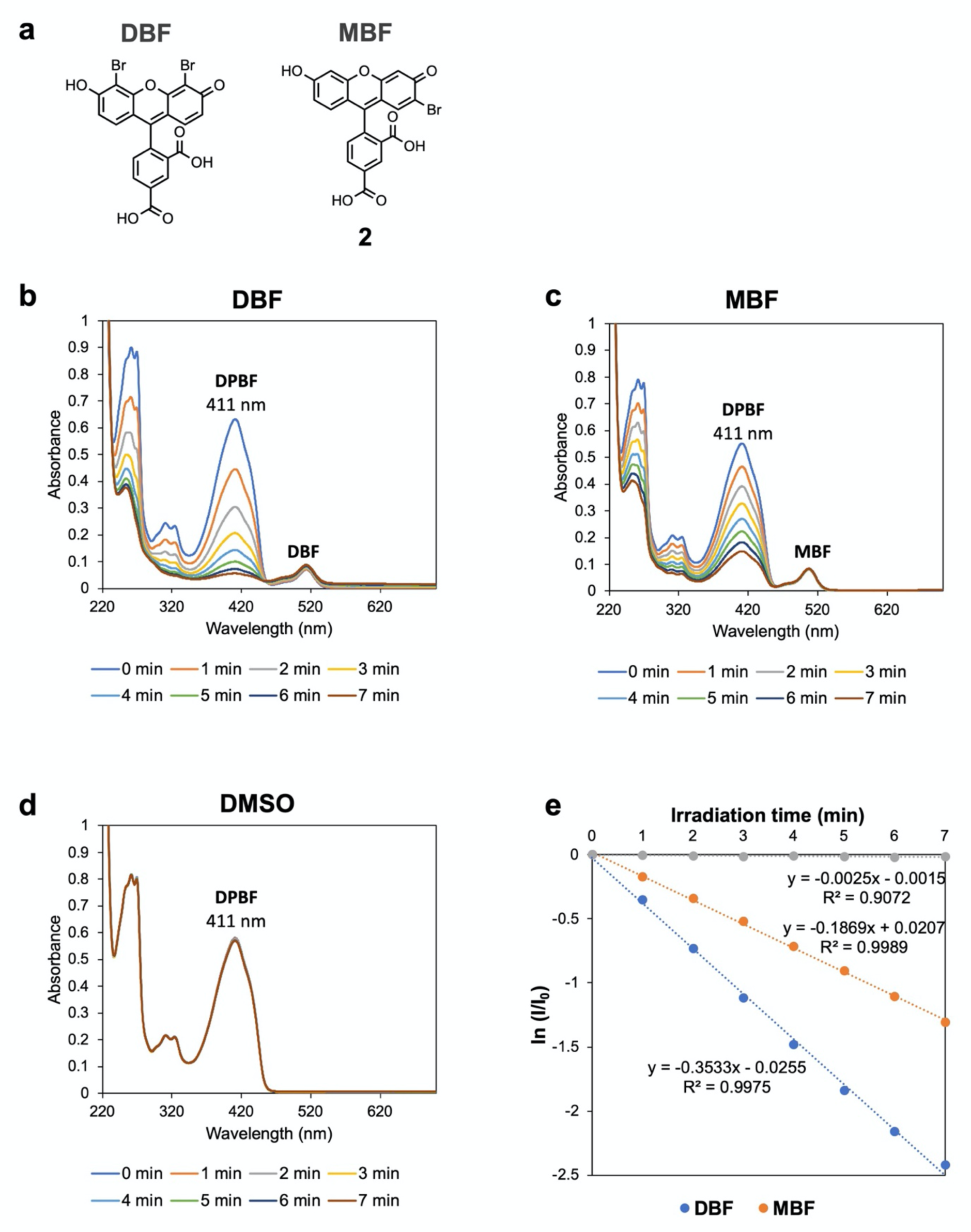
Estimation of the ^1^O_2_ quantum yield (Φ_Δ_) of MBF. The Φ_Δ_ of MBF was estimated by a relative spectrophotometric method using 1,3-diphenylisobenzofuran (DPBF) as the chemical trap for ^1^O_2_ and DBF as the reference photosensitiser. **(a)** Chemical structures of the DBF and MBF derivatives used. **(b–d)** Raw absorption spectra of showing the photoirradiation-dependent decrease in absorbance of DPBF in the presence of DBF and MBF but not DMSO. **(e)** Logarithm of DBPF concentration in the presence of MBF or DBF plotted against irradiation time. Φ_Δ_ of MBF was calculated from the slopes in this plot (see Experimental Section).

**Extended Data Fig. 2.**
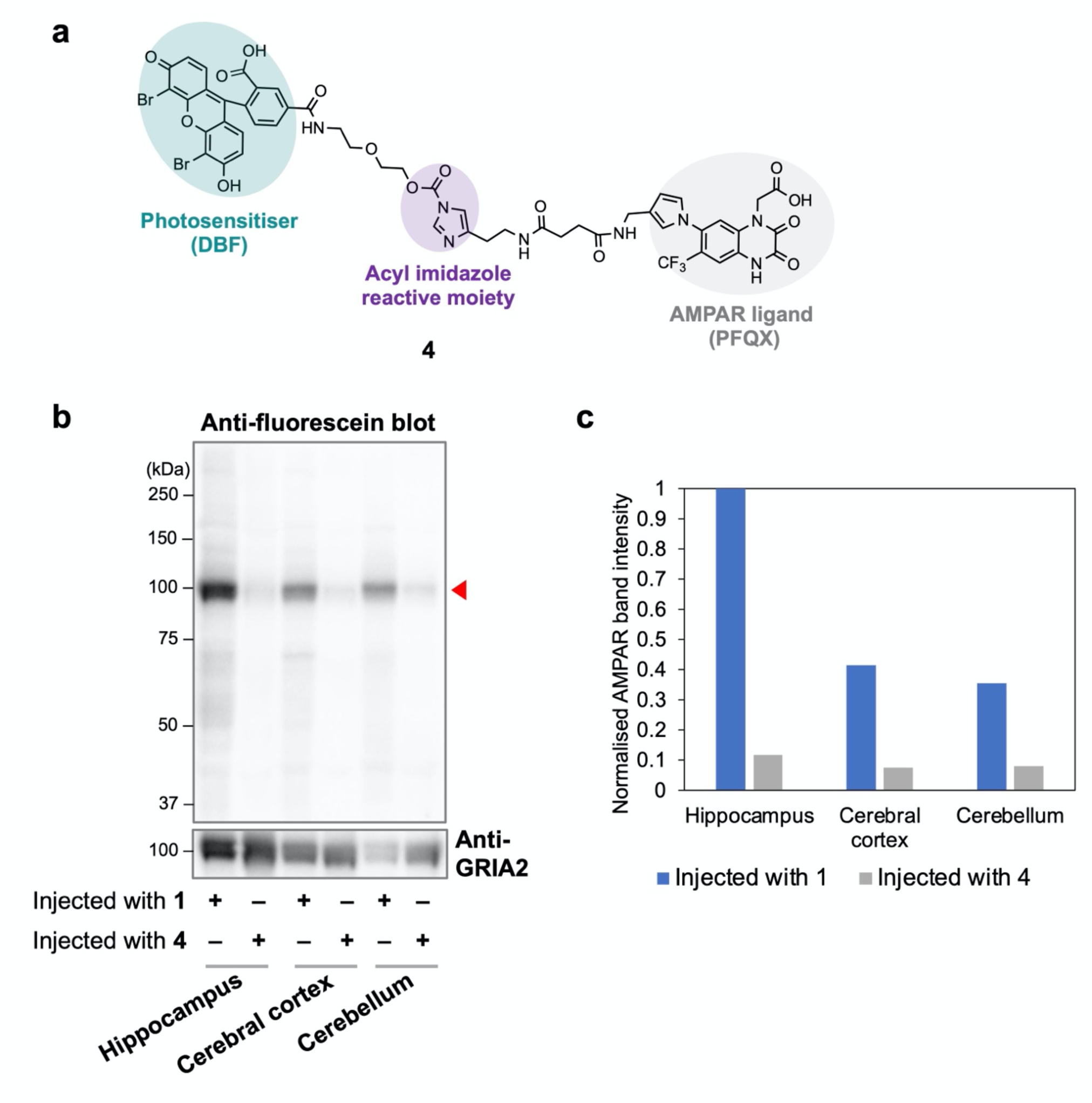
Comparison of the in vivo anchoring efficiency of MBF- and DBF-bearing LDAI reagents. **(a)** Chemical structure of **4**, an AMPAR-targeted LDAI reagent carrying DBF. **(b)** Western blot analysis of MBF and DBF anchoring onto AMPARs in brain tissue isolated from mice injected to the lateral ventricles with **1** or **4**. Red triangle points to the band corresponding to MBF- or DBF-modified AMPAR subunits. **(c)** Quantification of the ∼100 kDa band indicated by the red triangle in (b).

**Extended Data Fig. 3.**
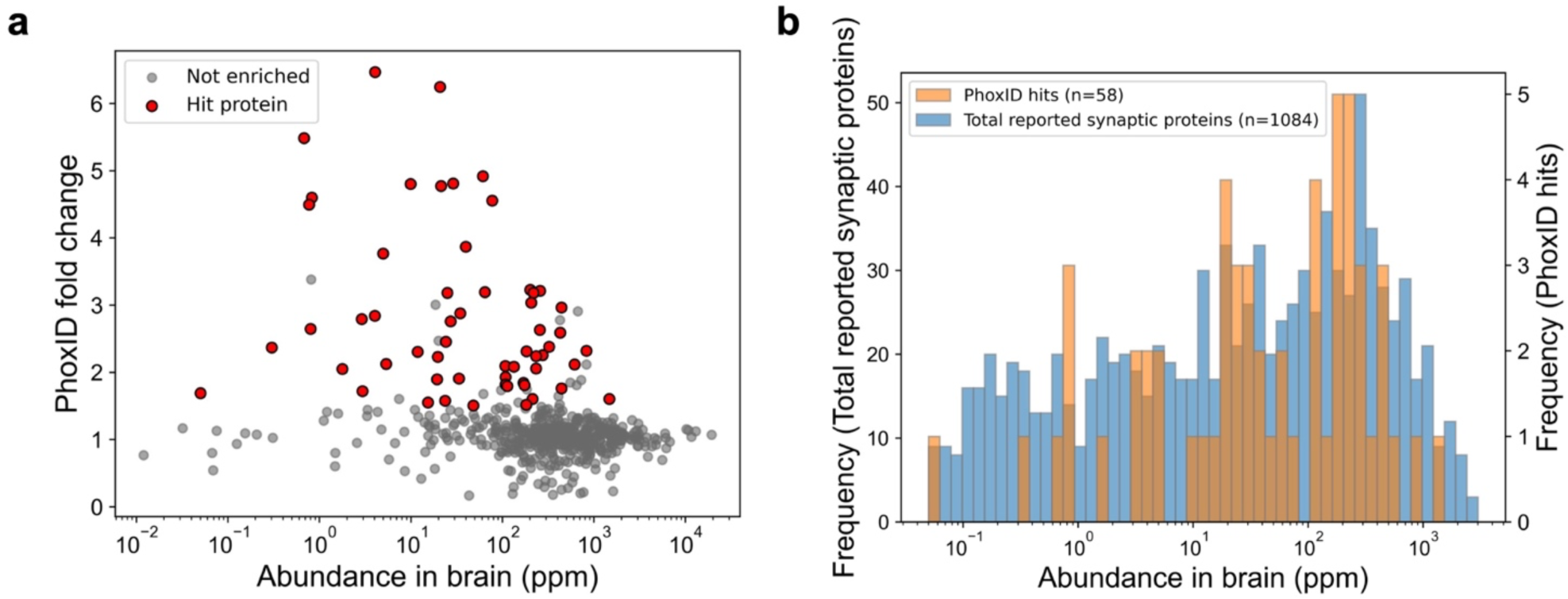
PhoxID can label proteins across a wide range of abundances. **(a)** Histogram showing the distribution of protein abundances among hippocampal AMPAR-proximal proteins as identified by PhoxID (orange) and the full list of reported synaptic proteins (blue, from SynGO release 20210225). Protein abundance data was retrieved from PAXdb (retrieved 2022-4). **(b)** PhoxID fold change of the hippocampal AMPAR-proximal hits plotted against their abundance in the brain.

**Extended Data Fig. 4.**
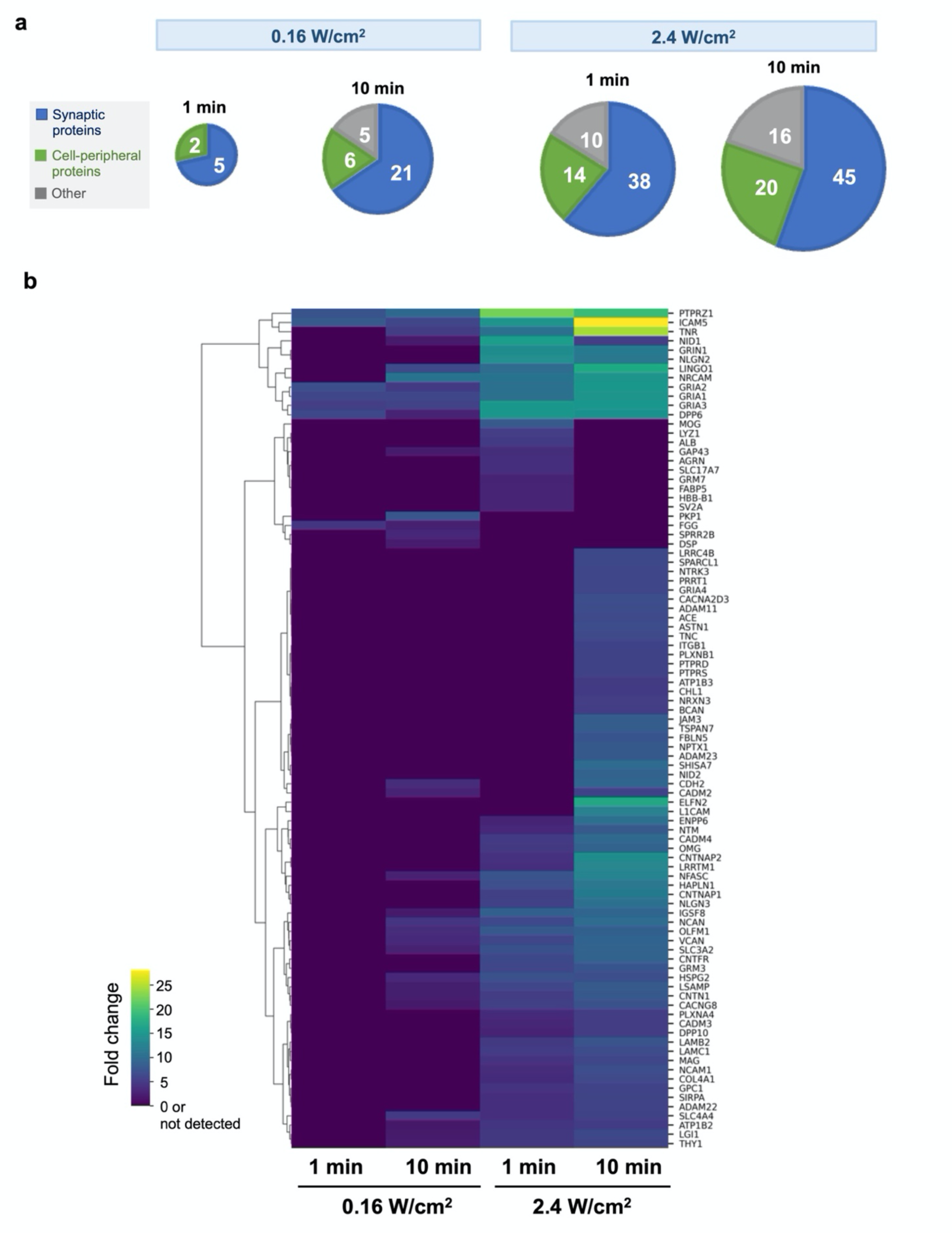
Supplementary data for Fig. 2h. **(a)** Fraction of hit proteins under each photoirradiation condition that are annotated to select synaptic GOCC terms and cell periphery (GO:0071944). Proteins that are annotated to both synaptic terms and the cell periphery are classified here as synaptic proteins. **(b)** Heatmap showing the fold change of proteins under various photoirradiation conditions. Data is only shown for proteins that met our hit protein criteria in at least one of the conditions tested. Mitochondrial/nuclear contaminant proteins are omitted.

**Extended Data Fig. 5.**
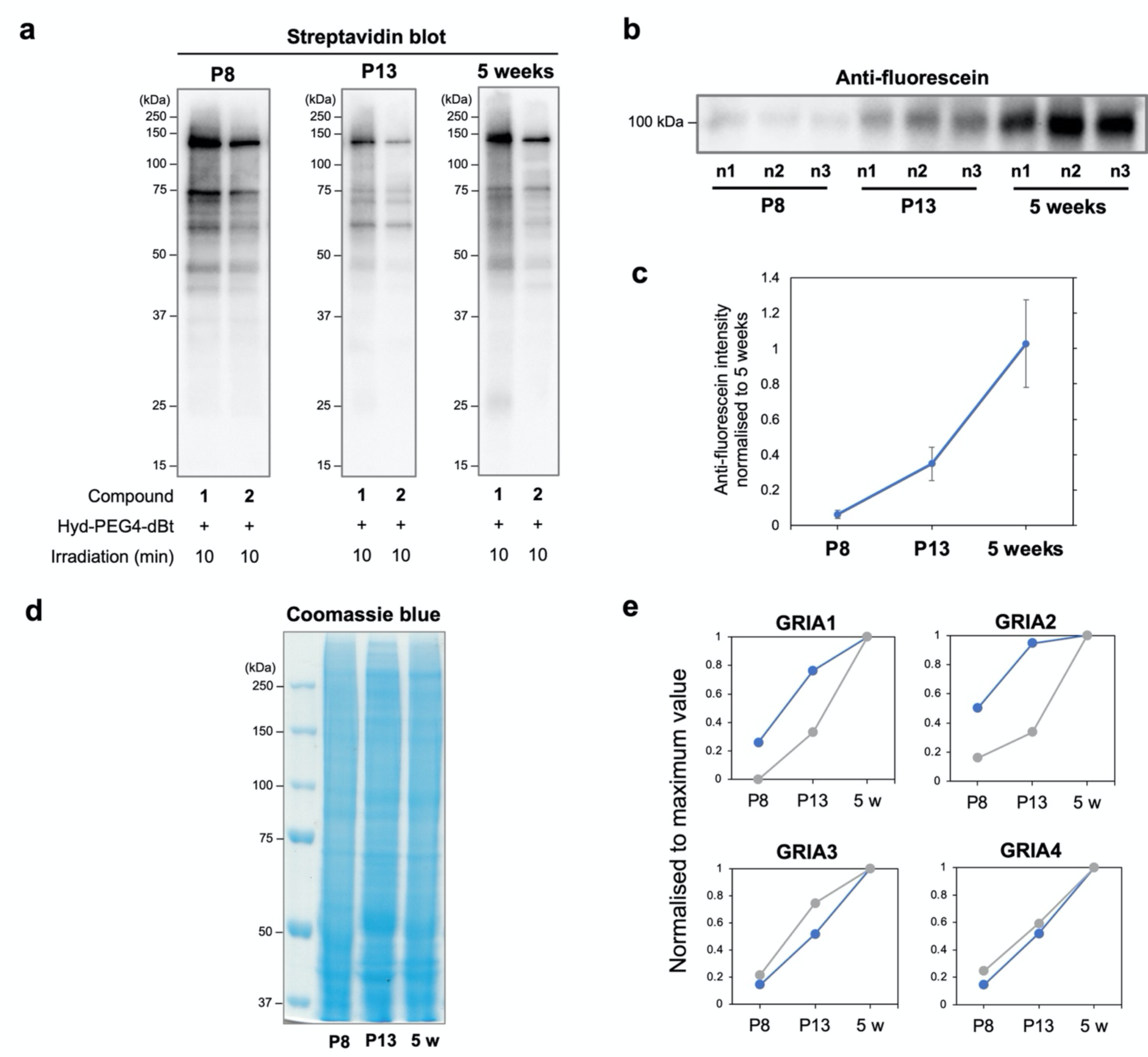
Supplementary data for Fig. 5. **(a)** Western blot analysis of MBF anchoring in the mouse cerebellum isolated from P8, P13, and 5-week-old mice injected with **1.** n1, n2, and n3 denote biological replicates. **(b)** Mean intensity of the western blot bands in (a). Data are normalised to the mean band intensity in 5-week-old mice. Error bars represent s.d. **(c)** Streptavidin blot of proteins labelled by PhoxID in the P8, P13, and 5-week-old cerebellum. Western blotting was performed after neutravidin enrichment. **(d)** Coomassie blue staining of the brain lysates analysed in Fig. 5e. **(e)** Comparison of the PhoxID fold change of the four AMPAR subunits at P8, P13, and 5 weeks (grey), and their protein expression levels as determined by western blot analysis in Fig. 5e (blue).

**Extended Data Fig. 6.**
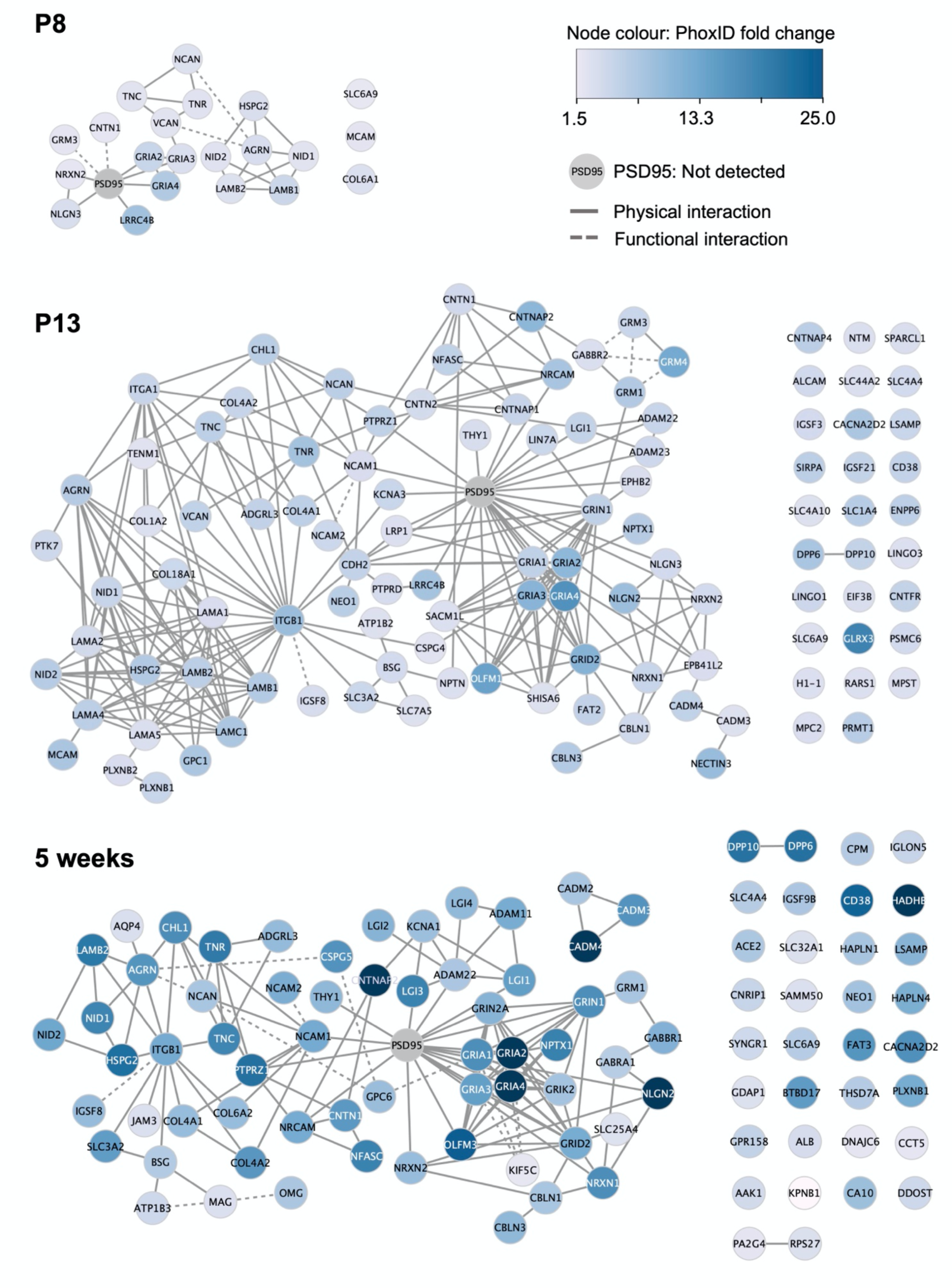
Network diagrams of known and predicted interactions between the PhoxID hit proteins at P8, P13, and week 5. Interaction information was obtained from STRING 11.5 (textmining and co-expression excluded) and several manual annotations (see Supplementary Table 5).

**Extended Data Fig. 7.**
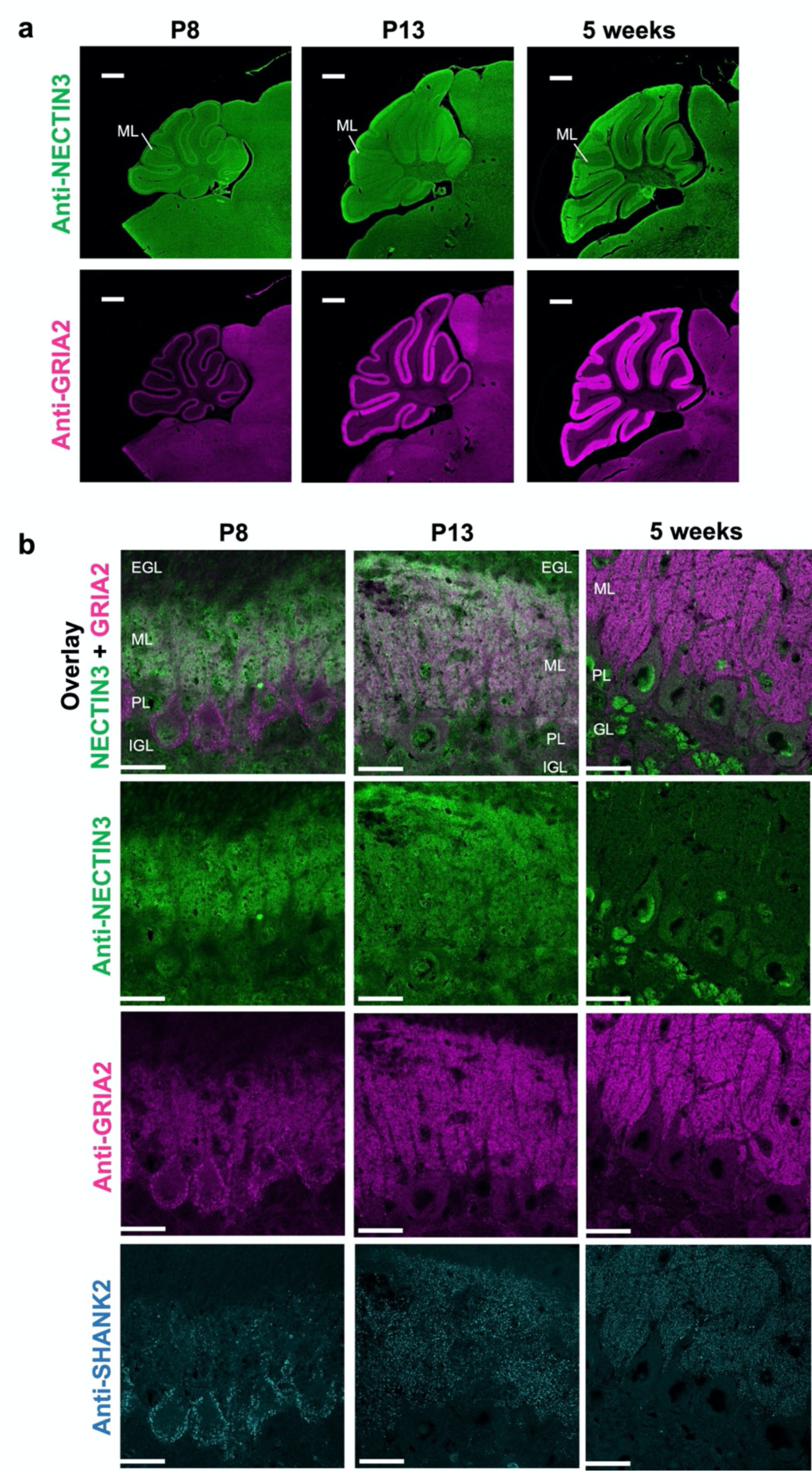
NECTIN3 is a developmentally regulated AMPAR-proximal protein. **(a)** Expression of NECTIN3 and AMPARs as detected by anti-NECTIN3 and anti-GRIA2 immunostaining on cerebellum sections from P8, P13, and 5-week-old mice. ML, molecular layer. Scale bar, 500 µm. **(b)** Magnified images of NECTIN3, GRIA2, and the postsynapse marker SHANK2 immunostained in the P8, P13, and 5-week-old cerebellum. Airyscan processing was applied. EGL, external granular layer. PL, Purkinje layer. IGL, internal granular layer. GL, granular layer. Scale bar, 20 µm.

**Extended Data Fig. 8.**
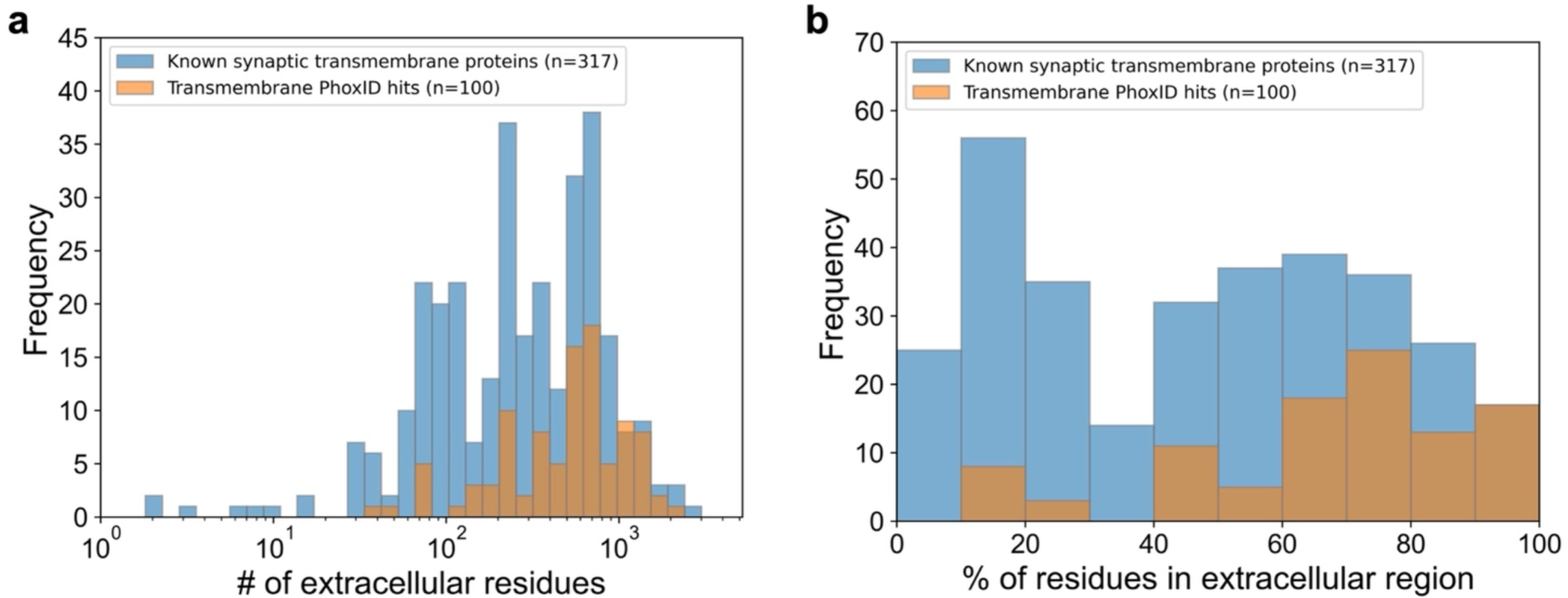
Proteins with large extracellular domains are more susceptible to labelling by PhoxID. Histograms showing the distribution of the total number of extracellular residues **(a)** and the percentage of the amino acid sequence in the extracellular region **(b)** among the hit proteins from all PhoxID experiments presented in this study (orange) and a list of reported transmembrane synaptic proteins (blue). Transmembrane synaptic proteins were defined as any protein in SynGO that has at least one extracellular topological domain annotated in UniProt (ver. 2023-2). PhoxID hit proteins were deemed transmembrane if they met the same criteria. The total number of extracellular residues for each protein was calculated by summing the number of residues in all extracellular domains recorded for the protein in UniProt. The percent of the sequence in the extracellular region was calculated for each protein by dividing the total number of amino acid residues in extracellular domains by the full length of the protein sequence then multiplying by 100.

**Extended Data Fig. 9.**
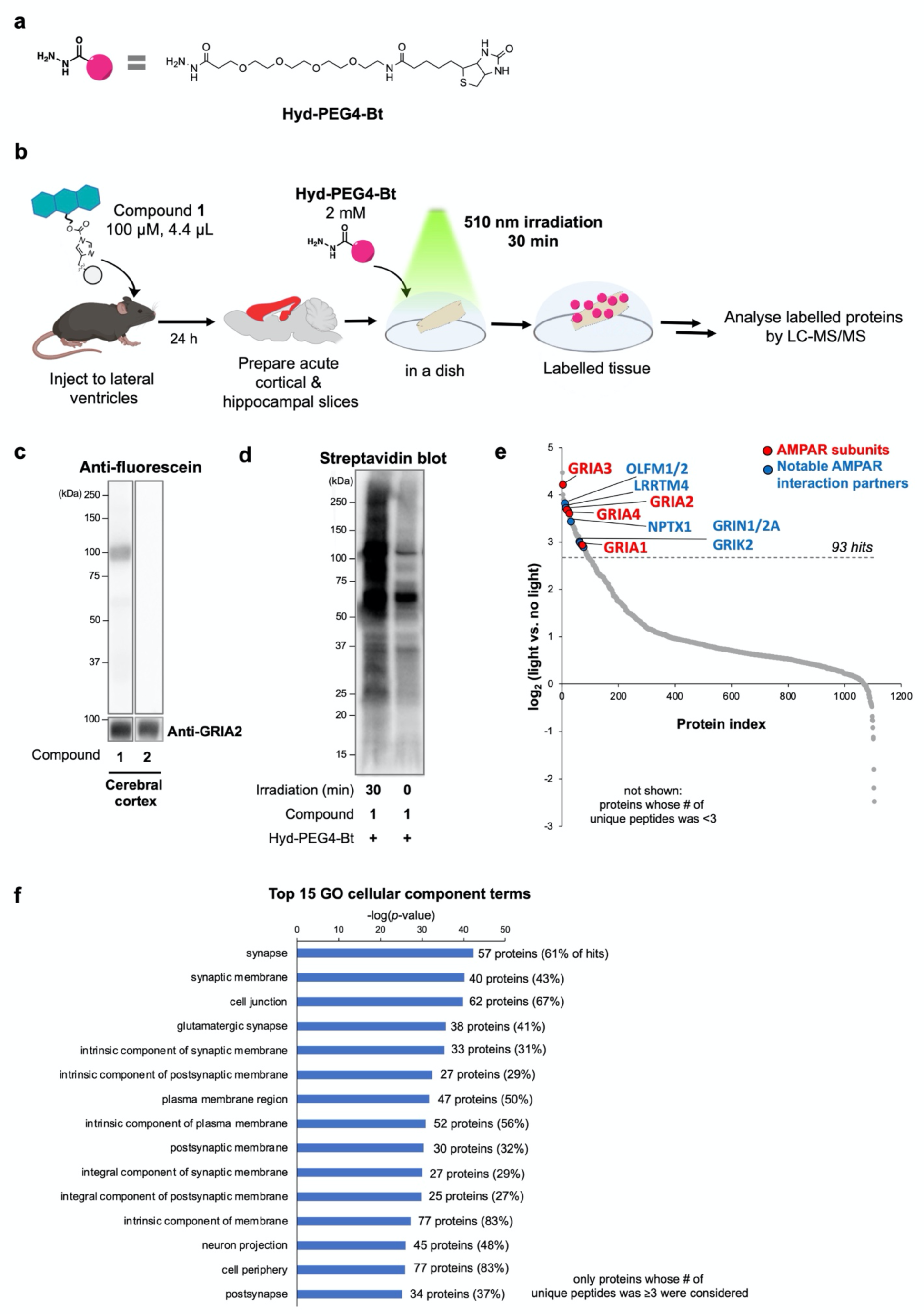
PhoxID can be performed in isolated brain slices. **(a)** Chemical structure of **Hyd-PEG4-Bt** bearing biotin as the enrichment handle. **(b)** Experimental workflow for PhoxID in acute brain slices. **(c)** Western blot analysis of MBF anchoring onto AMPARs in the cerebral cortex. **(d)** Streptavidin blot analysis of the proteins labelled by PhoxID in acute cortical and hippocampal slices. Western blotting was performed after neutravidin enrichment. **(e)** Fold change plot of the proteins identified by LC-MS/MS in a single biological replicate. **(f)** GOCC (complete) analysis of the PhoxID hits. The top 15 terms with the smallest raw *p*-value are shown. Percentages represent the fraction of hit proteins that are annotated to each GO term.

