## Supplementary Information for "Phototriggered profiling of receptor-proximal proteins in vivo in minutes"

### Supplementary Note

#### Definition of a "hit" protein

A "hit" protein was defined as any protein detected in a proteomics dataset that is retained after applying the following filters in sequence:

Filter 1: All detected proteins were initially filtered based on the number of unique peptides identified. For in vivo PhoxID in neonatal mice, only proteins whose unique peptide count was ≥2 were retained. For PhoxID in acute slices, only proteins detected with ≥3 unique peptides were retained. No filter based on unique peptide count was applied to other PhoxID data.

Filter 2: For proteomics experiments with biological replicates (n = 3), a filter based on fold change p-values was applied next. Only proteins whose fold change p-values (as calculated by ANOVA) was less than 0.1 (for AMPAR-targeted hippocampal and cerebellum PhoxID) or 0.05 (all other experiments) were retained. This filter was not applicable to proteomics datasets without biological replicates.

Filter 3: Finally, we defined contaminant proteins as those proteins annotated to the mitochondrion (GO:0005739) and/or the nucleus (GO: 0005634) but not the cell periphery (GO:0071944) in the Gene Ontology Cellular Component (GOCC) database (release 2022-03-10), and searched for the lowest fold change cut-off value for which the percent of the contaminant proteins among proteins above the value was below 10% (for PhoxID in neonatal mice) or 5% (all other experiments). This value was defined as the fold change cut-off, and proteins with fold changes at or above this value were classified as "hit" proteins.

### Experimental Section

#### General materials and methods for the biochemical/biological experiments.

Unless otherwise noted, all proteins/enzymes and reagents were obtained from commercial suppliers (Sigma-Aldrich, Tokyo Chemical Industry (TCI), Fujifilm Wako Pure Chemical Corporation, Bio-Rad, or Thermo Fisher Scientific) and used without further purification. SDS-

PAGE and western blotting were carried out using a Bio-Rad Mini-PROTEAN III electrophoresis apparatus. Samples were applied to SDS-PAGE and electrotransferred onto polyvinylidene fluoride membranes (Bio-Rad), followed by blocking with 5% nonfat dry milk in Tris-buffered saline containing 0.05% Tween 20. Primary antibodies are indicated in each experimental procedure, and anti-rabbit IgG-HRP conjugate (Cell Signaling Technology, 7074S, 1:4000), anti-mouse IgG-HRP conjugate (Cell Signaling Technology, 7076S, 1:4000), or anti-sheep IgG-HRP conjugate (Abcam, ab6747-1, 1:2000) were used as the secondary antibodies. Chemiluminescent signals generated with ECL Prime (GE Healthcare) were detected with a Fusion Solo S imaging system (Vilber Lourmat). InstantBlue Coomassie Protein Stain (Abcam, ab119211) was used to stain gels.

**General information for fluorescence imaging experiments.**

Confocal laser scanning microscopy (CLSM) was performed using TCS SP8 (Leica Microsystems, Germany) or LSM 800 (Carl Zeiss AG, Germany) equipped with a 5× objective (NA = 0.15 dry objective for TCS SP8), 5× objective (NA = 0.25 dry objective for LSM 800), 10× objective (NA = 0.40 dry objective), 63× objective (NA = 1.40 oil objective), and a GaAsP detector. The excitation laser was derived from a 488 nm, 561 nm, 640 nm diode laser (LSM 800) or a white laser (TCS SP8) and was set to an appropriate wavelength depending on the dye. LIGHTNING deconvolution process (LAS X 3.5.5, Leica Microsystems, Germany) or Airyscan mode (Carl Zeiss AG, Germany) was used for obtaining high-resolution images.

**Estimation of the $^1O_2$ quantum yield ($\Phi_\Delta$) of MBF.**

The $\Phi_\Delta$ of MBF was estimated by a relative spectrophotometric method using 1,3-diphenylisobenzofuran (DPBF) as the chemical trap for $^1O_2$ and DBF as the reference photosensitiser. A solution of MBF (1 μM, $\lambda_{max\ in\ EtOH\text{-}KOH}$ = 508 nm, $\varepsilon_{EtOH\text{-}KOH}$ = 106,000 M cm$^{-1}$)[S1] or DBF (1 μM, $\lambda_{max\ in\ EtOH\text{-}KOH}$ = 513 nm, $\varepsilon_{EtOH\text{-}KOH}$ = 94,100 M cm$^{-1}$, $\Phi_{\Delta,\ EtOH}$ = 0.29)[S1, S2] or DMSO and DPBF (30 μM) in 95% ethanol/0.01 M KOH solution in a quartz cuvette was illuminated with a homemade photoirradiation device ($\lambda_{max}$ = 512 nm, irradiance = 11–19 W/m$^2$, described previously[S3]). The absorption spectrum of the solution was measured at 0 min and after every minute of irradiation for 7 min using a Shimadzu UV-2600 spectrophotometer. The logarithm of DBPF concentration in the presence of MBF or DBF was plotted against irradiation time (**Extended Data Fig. 1e**), and $\Phi_{\Delta,\mathrm{MBF}}$ was calculated from the slopes of the two plots using the following equation:

$$\phi_{\Delta,MBF} = \phi_{\Delta,DBF} \frac{slope_{MBF}}{slope_{DBF}} \frac{(1 - 10^{-A_{DBF}})}{(1 - 10^{-A_{MBF}})}$$

where $A$ is the absorbance of DBF or MBF at 512 nm.

**Covalent anchoring of photosensitisers to neurotransmitter receptors in the mouse brain.** C57BL/6N mice (male, 5 weeks old, body weight = 18–23 g) were anaesthesised with a mixture of Domitor® (Nippon Zenyaku Kogyo Co., Ltd.), midazolam (Sandoz), and Vetorphale® (Meiji Seika Pharma Co., Ltd.) and a solution of **1**, **2**, **3**, or **4** (4.4 µL, 100 µM, phosphate-buffered saline (PBS (–)), 5% DMSO) was directly injected into both the left and right lateral ventricles (AP = -1.3 mm from bregma, ML = ±2 mm, depth = 2 mm) or the cerebellum (AP = -8 mm from bregma, ML = +1 mm) using a microinjector (Nanoliter 2010, World Precision Instruments) (600 nL/min). After 2 min, the injection capillary was withdrawn, and the anaesthesia antagonist Antisedan® (Nippon Zenyaku Kogyo Co. Ltd., 0.2 mL) was administered by intraperitoneal injection. After 24–28 h, the mouse was sacrificed under deep anaesthesia with isoflurane. The mouse brain tissue was isolated and lysed with 1% SDS RIPA buffer (pH 7.4, 25 mM Tris–HCl, 150 mM NaCl, 1% SDS, 1% Nonidet P-40, 0.25% deoxycholic acid) containing 1% protease inhibitor cocktail set III (Millipore, 539134). The protein concentrations of the supernatant were analysed by BCA assay (Pierce), and the normalised lysates were mixed with a quarter volume of 5 × sample buffer (pH 6.8, 312.5 mM Tris–HCl, 25% sucrose, 10% SDS, 0.025% bromophenol blue) containing 250 mM DTT and vortexed for 1 h at room temperature. SDS-PAGE and western blotting analyses were performed as described in “General methods for biochemical and biological experiments”. The MBF- or DBF-modified receptors were detected with an anti-fluorescein antibody (GeneTex,

GTX19491, 1:2000). Receptors modified with Alexa Fluor 647® were detected by in-gel fluorescence analysis following SDS-PAGE. Rabbit anti-Ionotropic Glutamate Receptor 2 [EPR18115] (Abcam, ab206293, 1:2000) and rabbit anti-GABA-A receptor gamma2 (Synaptic Systems, 224003, 1:2000) were used to detect GRIA2 and GABRG2 in mouse brain tissue, respectively.

**Perfusion fixation and brain slice preparation.**

Experiments were conducted according to a literature source.[S6] Mice injected with **1** or **3** according to "Covalent anchoring of photosensitisers to neurotransmitter receptors in the mouse brain" were transcardially perfused with ice-cold 4% paraformaldehyde (PFA)/PBS(–) (pH 7.4) (60 mL for 5-week-old mice, 20 mL for P8 and P13 mice) under deep anaesthesia with isoflurane. The isolated mouse brain was further fixed with 4% PFA/PBS(–) at overnight at 4 ºC. After washing with PBS(–) (×3), the brain was sagittally sliced along the midline into halves and immersed in 30% sucrose/PBS(–) at 4 °C over 1–3 days. The brain tissues were embedded in optimal cutting temperature (OCT) compound, frozen, and sagittally sliced into 15, 20, or 50 µm-thick sections using a cryostat (Leica Biosystems, CM1950).

**Immunostaining.**

For MBF and GRIA2 staining, cryosectioned brain slices (15 µm thick) were activated with antigen retrieval reagent ImmunoSaver (FUJIFILM Wako) at 80ºC for 20 min, washed thrice with PBS(–), and blocked with 10% normal goat serum (NGS) in PBS(–) containing 0.1% Triton X-100 for 30 min at room temperature. The primary antibody reaction was conducted with mouse anti-Glutamate Receptor 2, extracellular, clone 6C4 (Merck Millipore, MAB397, 1:500) and rabbit anti-fluorescein (Abcam, ab19491, 1:300) in PBS(–) containing 0.1% Triton X-100 overnight at 4 ºC. The secondary antibody reaction was conducted with a goat anti-mouse IgG (H+L)-Alexa Fluor® 647 conjugate (Invitrogen, A21235, 1:200) and a goat anti-rabbit IgG Alexa Fluor® 555 conjugate (Abcam, ab150078, 1:200) in PBS(–) containing 0.1% Triton X-100 for 1 h at room temperature. For GABRA1 staining, cryosectioned brain slices (50 µm thick) were treated with 0.2 % pepsin (from porcine gastric mucosa, Sigma, P-7012) in a buffer (pH 4.0) at 37 °C for 15 min. Permeabilisation and blocking was performed in PBS(–) containing 2% bovine serum albumin (BSA), 2% NGS and 0.2% Triton X-100 at room temperature for 30 min. After washing thrice with PBS(–), the slices were treated with rabbit anti-GABAA Receptor α1 (Millipore, 06-868, 1:300) in PBS(–) containing 0.1% Triton X-100 at 4°C overnight. The secondary antibody reaction was conducted with a goat anti-rabbit IgG H&L (Alexa Fluor® 546) (Invitrogen, A11071, 1:1000) in PBS(–) containing 0.1% Triton X-100 for 1 h at room temperature. For NECTIN3, GRIA2, and SHANK2 co-staining, cryosectioned brain slices (20 µm thick) from C57BL/6N mice were activated by autoclaving at 120 ºC in citric acid buffer for 20 min. Permeabilisation and blocking was performed in 10% NGS in PBS(–) containing 0.1% Triton X-100 at room temperature for 30 min. After washing thrice with PBS(–), the slices were treated with rabbit anti-Nectin-3/PVRL3 (Proteintech, 11213-1-AP, 1:100), mouse anti-Glutamate Receptor 2, extracellular, clone 6C4 (Merck Millipore, MAB397, 1:500), and guinea pig anti-Shank2 (Synaptic Systems, 162 204, 1:500) in PBS(–) containing 1% BSA and 0.1% Triton X-100 overnight at 4 ºC. The secondary antibody reaction was performed with a goat anti-rabbit IgG-Alexa Fluor® 555 conjugate (Abcam, ab150078, 1:200), a goat anti-mouse IgG (H+L)-Alexa Fluor® 647 conjugate (Invitrogen, A21235, 1:200), and a goat anti-guinea pig IgG H&L Alexa Fluor® 405 conjugate (Abcam, ab175678, 1:200) in PBS(–) containing 0.1% Triton X-100 for 1 h at room temperature.

**Photoinduced proximity labelling in 5-week-old mice.**

Mice injected with **1**, **2**, or **3** according to the protocol described in "Covalent anchoring of photosensitisers to neurotransmitter receptors in the mouse brain" were anaesthesised 24–28 h later by intraperitoneal injection of a mixture of Domitor®, midazolam, and Vetorphale®. For photoinduced labelling in the hippocampus, **Hyd-PEG4-dBt** (4.4 µL, 5 mM, PBS (–), 5% DMSO) was injected at AP = -2.3 mm from bregma, ML = ±1.6 mm, depth = 1.4 mm using a microinjector at a rate of 600 nL/min. For photoinduced labelling in the cerebellum, the same labelling reagent solution was injected into the vermis of cerebellar lobules V-VIII (depth = 0.75 mm from surface). 10 min after the end of injection, the injection sites were irradiated for 1–10 min via an optical fibre (Doric Lenses, 520 nm laser, LDFLS-520/060, fibre cannula:

FOC-C-200.1.25-0.22-3.0). Irradiance was 0.16 W/cm$^2$ for all in vivo PhoxID experiments unless specified otherwise. 5 min after the end of irradiation, the mouse brain was isolated and submerged in ice-cold cutting solution for 5 min (120 mM choline chloride, 3 mM KCl, 8 mM $MgCl_2$, 1.25 mM $NaH_2PO_4$, 28 mM $NaHCO_3$, 22 mM glucose, 0.5 mM ascorbic acid). Then, the hippocampus and the cerebellar vermis were macroscopically dissected and the tissue was homogenised with a pestle in 400 µL of 1% SDS-RIPA buffer (pH 7.4, 25 mM Tris–HCl, 150 mM NaCl, 1% SDS, 1% Nonidet P-40, 0.25% deoxycholic acid) containing 1% protease inhibitor cocktail set III (Millipore, 539134). The lysate was further homogenised with an ultrasonic cell disruptor (Branson Ultrasonics, Sonifier SFX 250), mixed with a 10x volume of chilled acetone (proteomics-grade), and kept at -80 ºC overnight prior to enrichment. For the MBF anchoring and photoinduced labelling protocol applied to 5-week-old mice for comparison with neonatal proteomes, see "Covalent anchoring and photoinduced proximity labelling in neonatal mice" below.

**Covalent anchoring and photoinduced proximity labelling in neonatal mice.**

C57BL/6N mice (P7, P12, or 5 weeks old) were anaesthetised with 2–4% isoflurane and clamped in a stereotactic apparatus. An incision was made in the scalp to expose the skull, which was lightly punctured at AP = -8 mm from bregma, ML = +1 mm) using a 27G needle. A solution of **1** or **2** (4.4 µL, 100 µM, PBS (–), 5% DMSO) was directly injected at this site (depth = 0.75 mm from surface) using a microinjector (600 nL/min) with a finely pulled glass pipette. After 24–28 h, the mice were once again anaesthetised with 2–4% isoflurane and the skull was punctured with a 27G needle, this time at AP = -8 mm from bregma, ML = -1 mm. A solution of **Hyd-PEG4-dBt** (4.4 µL, 5 mM, PBS(–), 5% DMSO) was injected at this site (depth = 0.75 mm from surface). 10 min after the end of injection, the injection site was irradiated for 10 min via an optical fibre (520 nm laser, Doric Lenses, LDFLS-520/060, fibre cannula: FOC-C-200.1.25-0.22-3.0, 0.16 W/cm$^2$) inserted at a depth of 0.75 mm. 5 min after the end of photoirradiation, the mice were decapitated and their brains were isolated. The cerebellum tissue was sagittally sliced into halves along the midline using a scalpel, and the right half was homogenised in 1% SDS RIPA buffer containing 1% protease inhibitor cocktail set III using a pestle. The lysate was further homogenised with an ultrasonic cell disruptor, mixed with a 10x volume of chilled acetone, and kept at -80 ºC overnight prior to enrichment.

**Photoinduced proximity labelling in acute brain slices**

Sagittal cortical and hippocampal slices (250 µm thick) were prepared from a C57BL/6N mouse (5 weeks old, male) injected to the lateral ventricles with **1** as described in "Covalent anchoring of neurotransmitter receptors in the mouse brain" and kept alive for 23 h. Slices were immersed in a solution of **Hyd-PEG4-Bt** (2 mM) in ACSF (125 mM NaCl, 2.5 mM KCl, 2 mM $CaCl_2$, 1 mM $MgCl_2$, 26 mM $NaHCO_3$, 1.25 mM $NaH_2PO_4$, 10 mM glucose) for 30 min at room temperature in a 35-mm glass-bottom dish (IWAKI) under 95% $O_2$/5% $CO_2$. The dishes containing the slices were then placed on aluminum foil and illuminated for 30 min at room temperature using a homemade photoirradiation device (512 nm, irradiance = 11–19 $W/m^2$, described previously[S3]) or kept in darkness for the same duration. After photoirradiation, the slices were washed thrice with carboxygenated ACSF (1 mL) and once with PBS(–) on ice. For each condition, 5 slices were pooled and lysed together in 1% SDS RIPA buffer containing 1% protease inhibitor cocktail set III. The lysate was further homogenised with an ultrasonic cell disruptor, mixed with a 10x volume of chilled acetone (proteomics-grade), and kept at -80 ºC overnight prior to enrichment.

**Enrichment of labelled proteins**

Protein precipitates in acetone were collected by centrifugation (20030 g, 10 min, 4 °C), washed with 4 mL of chilled acetone, and air-dried for 10 min at room temperature. The precipitate was dissolved in 300-400 µL of 1% SDS RIPA buffer by sonication (Branson Ultrasonics, Sonifier SFX 250). The solution was diluted with an equal volume of 0.1% SDS RIPA buffer to reduce the SDS concentration to ~0.5% and further sonicated before heating at 45 °C for 45 min. Following centrifugation (13500 rpm, 10 min, 4 °C), the protein concentration of the supernatant was determined by BCA assay (Pierce). The solution was diluted to a protein concentration of 1.5 mg/mL by adding ~0.5% SDS RIPA buffer, and 0.7–2 mg protein was added to High Capacity NeutrAvidin Agarose beads (Pierce, 29202, 200 µL) pre-washed with

PBS(–) in a 1.5 mL Protein LoBind tube (Eppendorf). After rotating at room temperature for 2 h, the beads were centrifuged (3000 g, 1 min, 4 °C) and washed with RIPA buffer (×4). The beads were transferred to a fresh 1.5-mL Protein LoBind tube and further washed with RIPA buffer (×4). The beads were then mixed with RIPA buffer supplemented with 4 mM biotin and incubated at 37 °C for 1 h. The suspension was then applied to a Micro Bio-Spin Chromatography Column (Bio-Rad, 732-6204) to remove the beads. The filtrate was mixed with a quarter volume of 5 × sample buffer containing 250 mM DTT and heated at 65 °C for 5 min. A streptavidin-HRP conjugate (Invitrogen, S911, 1:4000) was used to detect the labelled proteins by western blotting.

**Sample preparation for LC-MS/MS**

Enriched protein samples were loaded to a 10% SDS-PAGE gel (Bio-Rad, Mini-PROTEAN TGX Gels) and resolved for ~1 cm. The gel containing protein samples was manually excised into dices and fixed with 45% methanol/water containing 5% acetic acid for 20 min. The fixed gel pieces were washed with 50% methanol aq. and pure water, then dehydrated with acetonitrile. The dehydrated gels were swelled with 200 µL of 10 mM DTT in 100 mM TEAB (triethylamine bicarbonate) buffer and heated at 56°C for 30 min. The DTT solutions were replaced with 55 mM iodoacetamide in 100 mM TEAB buffer, and the gels were incubated at 37°C in the dark for 30 min. The gels were then dehydrated in acetonitrile, rehydrated in 100 mM TEAB buffer, and dehydrated in acetonitrile again. The gel was swelled in 100 mM TEAB buffer containing 10 ng/µL Sequence Grade Trypsin (Promega, V5113) and incubated overnight at 37°C. An extraction solution (50% acetonitrile containing 0.1% TFA) was added to the gels and the gels were sonicated for 10 min in a bath sonicator. The supernatant was collected in a new Protein LoBind tube, and this process was repeated once with 50% acetonitrile containing 0.1% TFA and twice with 100% acetonitrile containing 0.1% TFA. The collected peptide solution was concentrated by a centrifugal concentrator. The extracted peptides were further purified by GL-Tip SDB (GL Sciences) according to the manufacturer's instructions.

**NanoLC–MS/MS analyses**

NanoLC–MS/MS analyses were performed on a Q Exactive mass spectrometer (Thermo Fisher Scientific) and an Ultimate 3000 nanoLC pump (AMR) as described previously.[S7] Samples were automatically injected using the PAL system (CTC analytics, Zwingen, Switzerland) into a peptide L-trap column OSD (5 μm) attached to an injector valve for desalinating and concentrating peptides. After washing the trap with MS-grade water containing 0.1% TFA and 2% acetonitrile, the peptides were loaded into a nano-HPLC capillary column (C18 packed with the gel particle size of 3 μm, 0.1 × 125 mm, Nikkyo Technos, Tokyo Japan) by switching the valve. The injection volume was 5 or 10 μL and the flow rate was 500 nL/min. The mobile phases consisted of (A) 0.5% acetic acid and (B) 0.5% acetic acid and 80% acetonitrile. A two-step linear gradient of 5–45% B in 60 min, 45–95% B in 1 min, 95% B for 20 min was employed. Spray voltages of 2,000 V were applied. The mass scan ranges were m/z 350–1,800, and top ten precursor ions were selected in each MS scan for subsequent MS/MS scans. The normalised collision energy was set to be 30. The raw MS data files were analysed by Proteome Discoverer 3.0 (Thermo Fisher Scientific) to create peak lists based on the recorded fragmentation spectra. Peptides and proteins were identified by means of automated database searching using Sequest HT and CHIMERYS (Thermo Fisher Scientific) against Mus musculus, UniProtKB/SWISS-PROT (release 2022-12) with a precursor mass tolerance of 10 p.p.m., a fragment ion mass tolerance of 0.02 Da, and trypsin specificity that allows for up to two missed cleavages. A reversed decoy database search was conducted with Percolator node, setting the false discovery rate (FDR) threshold to 5% at the peptide level. We performed technical duplicates of three independent biological replicates in label-free quantification analysis. Protein abundances were normalised across samples on the abundances of three known endogenously biotinylated proteins (UniProt IDs: Q99MR8, Q05920, Q91ZA3) and missing values were replaced with random values sampled from the lowest 5% of all detected values. Common mass spectrometry contaminant proteins, namely those found in the CRAPome database[S8] modified by Thermo Fisher Scientific, were removed from the protein list prior to further data analysis.

**Western blot analysis of protein expression levels at different postnatal ages.**

Brains were isolated from C57BL/6N mice at postnatal day 8, day 13, and week 5 under anaesthesia with isoflurane and immersed in ice-cold cutting solution for 5 min. The cerebellum was sagittally sliced along the midline and the right half was lysed in 300–400 µL 1% SDS RIPA buffer containing 1% protease inhibitor cocktail III using a pestle and an ultrasonic cell disrupter (Branson Ultrasonics, Sonifier SFX 250). After rotating for 30 min at 4 ºC, the lysate was centrifuged (13500 rpm, 10 min, 4 ºC) and the protein concentration of the supernatant was determined by BCA assay. Then, the protein concentration was normalised across different postnatal ages. For detection of LRRC4B, NECTIN3, and GRIA1–4, a quarter volume of 5x sample buffer containing 250 mM DTT was added to the protein extract. For detection of IGSF3, a quarter volume of 5x sample buffer without 250 mM DTT was added. After vortexing for 1 h at room temperature, the samples were analysed by western blotting. The primary antibody reactions were performed with rabbit anti-LRRC4B/NGL3 (Bioss, bs-11090R, 1:1000), rabbit anti-Nectin-3/PVRL3 (Proteintech, 11213-1-AP, 1:1000), sheep anti-IGSF3 (R&D Systems, AF4788, 0.5 µg/mL), rabbit anti-Glutamate Receptor 1 (AMPA subtype) [EPR5479] (Abcam, ab109450, 1:2000) (GRIA1), rabbit anti-Ionotropic Glutamate Receptor 2 [EPR18115] (Abcam, ab206293, 1:2000) (GRIA2), rabbit anti-AMPA Receptor 3 (GluA3) (D47E3) (Cell Signaling Technology, 4676T, 1:2000) (GRIA3), and rabbit anti-AMPA Receptor 4 (GluA4) (D41A11) XP® (Cell Signaling Technology, 8070S, 1:1000) (GRIA4).

**Data visualisation**

Network diagrams were created using Cytoscape (v. 3.9.1). Principal component analysis was conducted in Python using the scikit-learn library. All other plots were created in Microsoft Excel or in Python using the matplotlib and seaborn libraries. Cartoon figures were created using BioRender (https://www.biorender.com/) and Microsoft PowerPoint.

**General materials and methods for organic synthesis**

All chemical reagents and solvents were obtained from commercial suppliers (Tokyo Chemical Industry (TCI), Sigma-Aldrich, Thermo Fisher Scientific, Fujifilm-Wako Pure Chemical Corporation, Watanabe Chemical Industries, or Kanto Chemical Co., Inc.) and used without further purification. Thin layer chromatography (TLC) was performed on silica gel 60 F254 precoated aluminum sheets (Merck) and visualised by fluorescence quenching, fluorescence by 365 nm excitation, and ninhydrin staining. Chromatographic purification was accomplished using flash column chromatography on silica gel 60 N (neutral, 40–50 μm, Kanto Chemical). $^{1}$H- and $^{13}$C- NMR spectra were recorded in deuterated solvents on a Varian Mercury 400 (400 MHz) spectrometer or JEOL JNM-ECZ600R/S1 (600 MHz), and calibrated to tetramethylsilane (= 0 ppm) or residual solvent peak. Multiplicities are abbreviated as follows: s = singlet, brs= broad singlet, d = doublet, t = triplet, q = quartet, m = multiplet, dd = double doublet. High-resolution mass spectra were measured on an Exactive Plus (Thermo Fisher Scientific) equipped with electron spray ionisation (ESI). Reversed-phase HPLC (RP-HPLC) was carried out on a Hitachi Chromaster system equipped with a 5410 UV detector and a 5420 UV-Vis detector.

### Synthesis

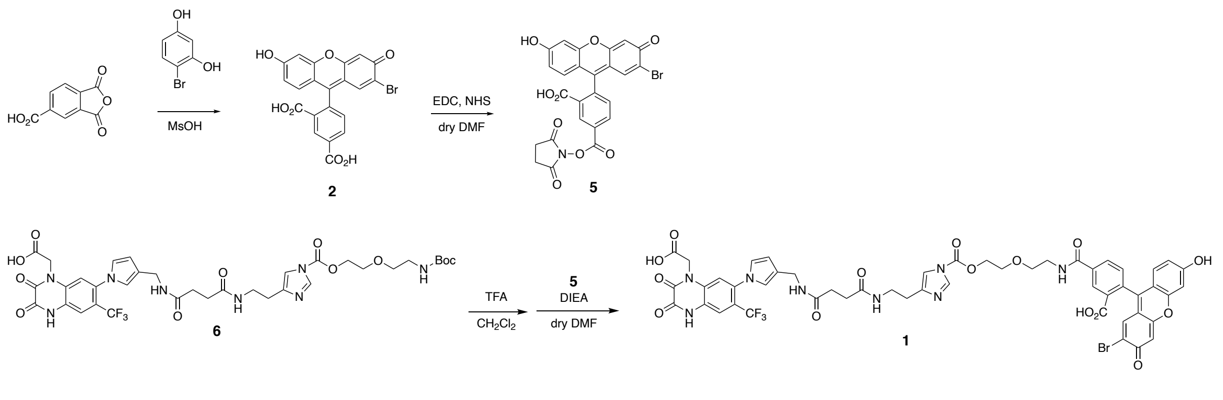


**Scheme 1** Synthetic scheme of **1**

*Compound* ***2***

To a stirred solution of trimellitic anhydride (96 mg, 0.50 mmol) in methanesulfonic acid (MsOH, 2.0 mL) was added 4-bromoresorcinol (189 mg, 1.0 mmol). The mixture was stirred at 50 °C for 30 h and then poured into water (20 mL). The solution was extracted with AcOEt (20 mL), and the organic layer was washed with water (10 mL x 4) and dried over $MgSO_4$. The solvent was removed under reduced pressure, and the residue was purified by flash chromatography on silica gel ($CHCl_3$ : MeOH : AcOH = 95 : 4 : 1) and RP-HPLC (0.1% trifluoroacetic acid (TFA)-AcCN : 0.1% TFA-$H_2O$ = 10 : 90 (0 min), 10 : 90 (5 min), 34 : 66 (50 min)) with a YMC-Pack ODS-A column (5 μm, 250 × 20 mm) at a flow rate of 9.9 mL/min. The target fraction was lyophilised to yield 2'-bromo-5-carboxyfluorescein (**2**) (35 mg, 77 μmol, 15%, single isomer) as an orange solid.

$^1$H NMR (600 MHz; $CD_3OD$): δ 8.63 (s, 1H), 8.41 (d, 1H, $J$ = 7.8 Hz), 7.34 (d, 1H, $J$ = 7.8 Hz), 6.86 (s, 1H), 6.84 (s, 1H), 6.72 (d, 1H, $J$ = 1.8 Hz), 6.40 (d, 1H, $J$ = 8.4 Hz), 6.58 (dd, 1H, $J$ = 1.8, 8.4 Hz).

HRMS (ESI): Calcd for $(C_{21}H_{10}Br_1O_7)^-$ $[M–H]^-$: 452.9615, found: 452.9618.

*Compound* ***5***

To a stirred solution of **2** (19 mg, 42 μmol) in dry *N,N*-dimethylformamide (DMF, 1.0 mL) was added 1-ethyl-3-(3-dimethylaminopropyl)carbodiimide hydrochloride (EDC, 9.6 mg, 50 μmol) and *N*-hydroxysuccinimide (NHS, 4.8 mg, 42 μmol). The mixture was stirred at room temperature for 14 h and purified by RP-HPLC (0.1% TFA-AcCN : 0.1% TFA-$H_2O$ = 35 : 65 (0 min), 60 : 40 (25 min)) with a YMC-Pack ODS-A column (5 μm, 250 × 20 mm) at a flow rate of 9.9 mL/min. The target fraction was lyophilised to yield **5** (13 mg, 23 μmol, 54%) as an orange solid.

$^1$H NMR (400 MHz; $CD_3CN$): δ 8.68 (d, 1H, $J$ = 1.6 Hz), 8.46 (dd, 1H, $J$ = 1.6, 8.4 Hz), 7.47 (d, 1H, $J$ = 8.4 Hz), 7.01 (s, 1H), 6.91 (s, 1H), 6.75 (d, 1H, $J$ = 2.4 Hz), 6.70 (d, 1H, $J$ = 8.8 Hz), 6.59 (dd, 1H, $J$ = 2.4, 8.8 Hz), 2.91 (s, 4H).

*Compound **1***

To a stirred solution of **6**[S4] (9.0 mg, 11 μmol) in dichloromethane (1.0 mL) was added TFA (500 μL). The mixture was stirred at room temperature for 1 h. TFA was removed by co-evaporation with toluene (500 μL) twice. To a stirred solution of this residue in dry DMF (500 μL) was added **5** (7.4 mg, 13 μmol) and *N,N*-diisopropylethylamine (DIEA, 14 μL, 80 μmol). The mixture was stirred at room temperature for 1.5 h and purified by RP-HPLC (10 mM triethylammonium acetate (TEAA)-AcCN : 10 mM TEAA-$H_2O$ = 10 : 90 (0 min), 10 : 90 (5 min), 40 : 60 (35 min)) with a YMC-Pack ODS-A column (5 μm, 250 × 20 mm) at a flow rate of 9.9 mL/min. The target fraction was lyophilised to yield **1** (10 mg, 7.4 μmol, 66%) as an orange solid.

$^1$H NMR (600 MHz; $CD_3OD$): δ 8.48 (s, 1H), 8.18 (s, 1H), 8.04 (dd, 1H, *J* = 1.2, 7.8 Hz), 7.54 (s, 1H), 7.35 (s, 1H), 7.32 (d, 1H, *J* = 7.8 Hz), 7.19 (s, 1H), 7.12 (s, 1H), 6.92 (d, 1H, *J* = 9.0 Hz), 6.79–6.75 (m, 2H), 6.72 (d, 1H, *J* = 2.4 Hz), 6.65–6.62 (m, 2H), 6.18 (s, 1H), 4.72 (brs, 2H), 4.57 (t, 2H, *J* = 4.8 Hz), 4.23 (s, 2H), 3.87 (t, 2H, *J* = 4.8 Hz), 3.75 (t, 2H, *J* = 5.4 Hz), 3.63 (t, 2H, *J* = 5.4 Hz), 3.37 (t, 2H, *J* = 6.0 Hz), 2.66 (t, 2H, *J* = 6.0 Hz), 2.48–2.45 (m, 4H).

$^{13}$C NMR (150 MHz; $CD_3OD$): δ 177.95, 174.64, 174.18, 171.75, 171.66, 157.78, 157.41, 156.87, 155.88, 149.84, 142.03, 138.48, 137.39, 135.96, 133.40, 131.64, 131.31, 129.42, 128.41, 126.49, 125.32, 125.10, 123.47, 123.38, 122.89, 122.47, 122.26, 118.20, 118.15, 117.74, 115.42, 115.38, 114.56, 113.38, 110.45, 104.49, 103.72, 70.54, 69.48, 68.51, 41.02, 39.69, 37.22, 32.49, 32.43, 28.70

HRMS (ESI): Calcd for $(C_{51}H_{43}Br_1F_3N_8O_{15})^+$ $[M+H]^+$: 1143.1978, found: 1143.1959.

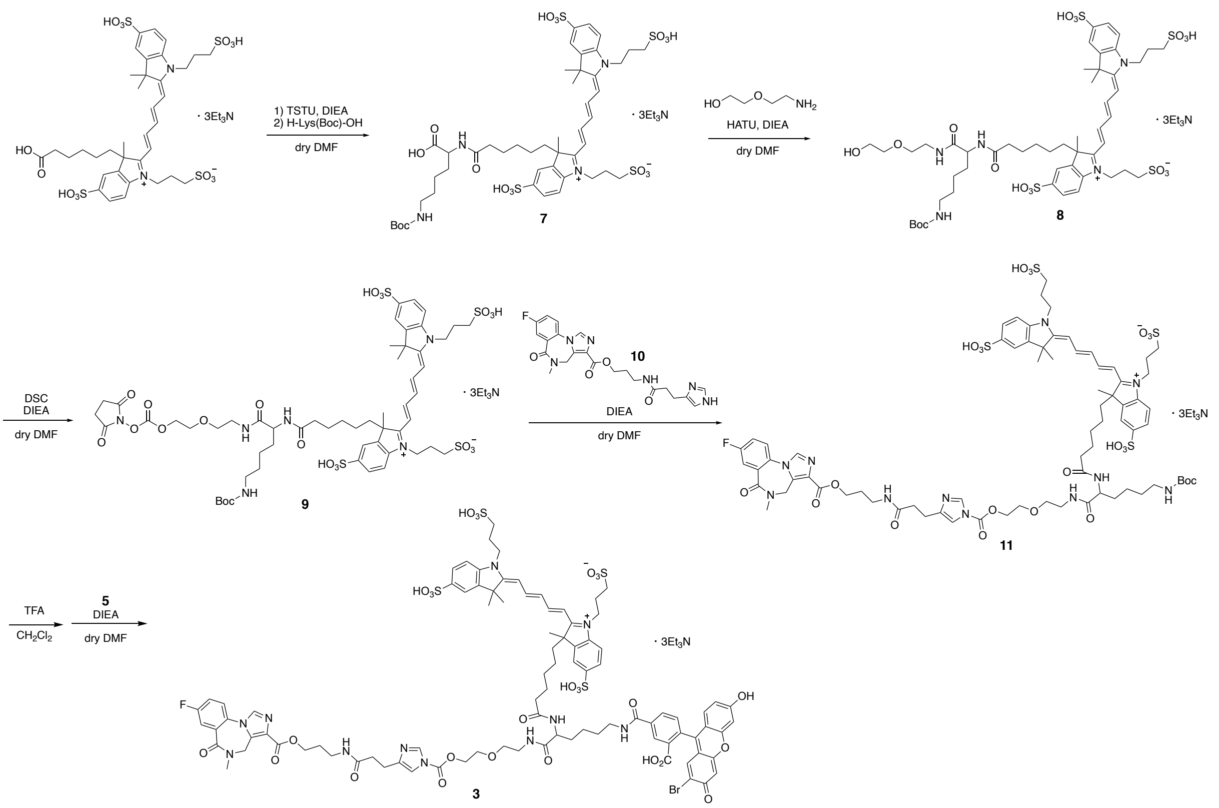

**Scheme 2** Synthetic scheme of **3**

*Compound 7*

To a stirred solution of Alexa Fluor$^{TM}$ 647 carboxylic acid tris (triethylammonium) salt (5.0 mg, 4.3 μmol) in dry DMF (500 μL) was added *N,N,N',N'*-tetramethyl-*O*-(*N*-succinimidyl)uranium tetrafluoroborate (TSTU, 1.3 mg, 4.3 μmol) and DIEA (4.5 μL, 26 μmol). After stirring for 2 h at room temperature, H-Lys(Boc)-OH (1.06 mg, 4.30 μmol) was added to the solution. After 2 h, the solvent was removed under reduced pressure, and the residue was purified by RP-HPLC (10 mM TEAA-AcCN : 10 mM TEAA-$H_2O$ = 5 : 95 (0 min), 20 : 80 (30 min)) with a YMC-Pack Pro C18 RS column (5 μm, 250 × 10 mm) at a flow rate of 3.0 mL/min. The target fraction was lyophilised to yield **7** (6.0 mg, 4.3 μmol, quant) as a blue film.
HRMS (ESI): Calcd for $(C_{47}H_{64}N_4O_{17}S_4)^{2-}$ $[M–2H]^{2–}$: 542.1585, found: 542.1580.

*Compound **8***

To a stirred solution of **7** (5.0 mg, 3.6 μmol) in dry DMF (500 μL) was added 1-[bis(dimethylamino)methylene]-1*H*-1,2,3-triazolo[4,5-*b*]pyridinium 3-oxide hexafluoro-phosphate (HATU, 1.6 mg, 4.2 μmol), 2-(2-aminoethoxy)ethanol (0.75 mg, 7.2 μmol) and DIEA (3.1 μL, 18 μmol). The mixture was stirred at room temperature for 5 h. The solvent was removed under reduced pressure, and the residue was purified by RP-HPLC (10 mM TEAA-AcCN : 10 mM TEAA-$H_2O$ = 5 : 95 (0 min), 25 : 75 (20 min)) with a YMC-Pack Pro C18 RS column (5 μm, 250 × 10 mm) at a flow rate of 3.0 mL/min. The target fraction was lyophilised

to yield **8** (5.1 mg, 3.4 µmol, 94%) as a blue film.
HRMS (ESI): Calcd for $(C_{51}H_{73}N_5O_{18}S_4)^{2-}$ $[M–2H]^{2-}$: 542.1585, found: 542.1580.

*Compound* ***9***

To a stirred solution of **8** (5.1 mg, 3.4 µmol) in dry DMF (500 µL) was added di(*N*-succimidyl)carbonate (DSC, 30 mg, 117 µmol) and DIEA (28 µL, 160 µmol). The mixture was stirred at room temperature for 18 h. The solvent was removed under reduced pressure, and the residue was purified by RP-HPLC (10 mM TEAA-AcCN : 10 mM TEAA-$H_2O$ = 5 : 95 (0 min), 30 : 70 (25 min)) with a YMC-Pack Pro C18 RS column (5 µm, 250 × 10 mm) at a flow rate of 3.0 mL/min. The target fraction was lyophilised to yield **9** (3.0 mg, 1.9 µmol, 53%) as a blue film.
HRMS (ESI): Calcd for $(C_{56}H_{76}N_6O_{22}S_4)^{2-}$ $[M–2H]^{2-}$: 656.1953, found: 656.1946.

*Compound* ***11***

To a stirred solution of **9** (3.0 mg, 1.9 µmol) in dry DMF (300 µL) was added **10**[S9] (4.2 mg, 9.3 µmol) and DIEA (3.2 µL, 19 µmol). The mixture was stirred at room temperature for 6 h. The solvent was removed under reduced pressure, and the residue was purified by RP-HPLC (10 mM TEAA-AcCN : 10 mM TEAA-$H_2O$ = 10 : 90 (0 min), 35 : 65 (25 min)) with a YMC-Pack Pro C18 RS column (5 µm, 250 × 10 mm) at a flow rate of 3.0 mL/min. The target fraction was lyophilised to yield **11** (1.0 mg, 0.51 µmol, 28%) as a blue film.
HRMS (ESI): Calcd for $(C_{74}H_{94}F_1N_{11}O_{23}S_4)^{2-}$ $[M–2H]^{2-}$: 825.7701, found: 825.7709.

*Compound* ***3***

To a stirred solution of **11** (1.0 mg, 0.51 µmol) in dichloromethane (1.0 mL) was added TFA (300 µL). The mixture was stirred at room temperature for 0.5 h. TFA was removed by co-evaporation with toluene (500 µL) three times. To a stirred solution of this residue in dry DMF (500 µL) was added **5** (1.4 mg, 2.6 µmol) and DIEA (2.7 µL, 15 µmol). The mixture was stirred at room temperature for 3 h. The solvent was removed under reduced pressure, and the residue was purified by RP-HPLC (10 mM TEAA-AcCN : 10 mM TEAA-$H_2O$ = 10 : 90 (0 min), 23 : 77 (26 min)) with a YMC-Pack Pro C18 RS column (5 µm, 250 × 10 mm) at a flow rate of 3.0 mL/min. The target fraction was lyophilised to yield **3** (0.79 mg, 0.35 µmol, 68%) as a blue film.
HRMS (ESI): Calcd for $(C_{90}H_{95}Br_1F_1N_{11}O_{27}S_4)^{2-}$ $[M–2H]^{2-}$: 993.7230, found: 993.7234.

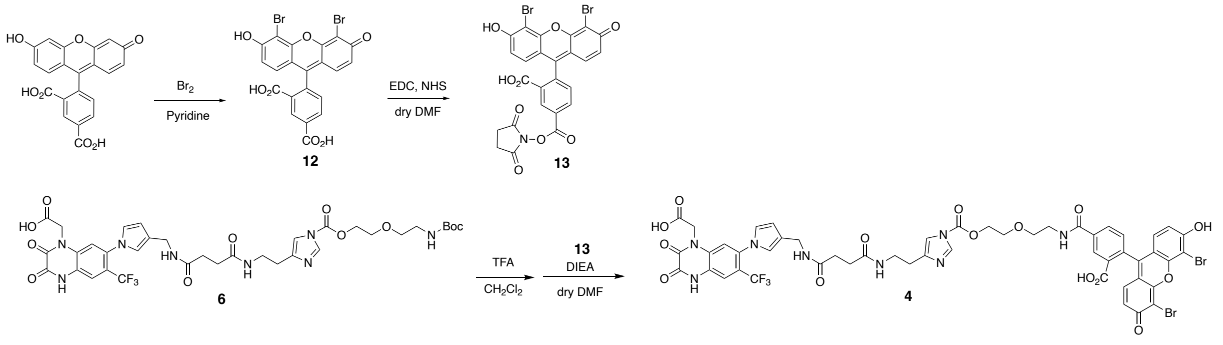


**Scheme 3** Synthetic scheme of **4**

*Compound **12***

To a stirred solution of 5-carboxyfluorescein (225.8 mg, 0.600 mmol) in pyridine (1.5 mL) was dropwise added bromine (61.2 μL, 1.19 mmol) in pyridine (1.5 mL) at 0 °C. The mixture was warmed at room temperature and allowed to stir for 3 h. The reaction mixture was diluted by 1N HCl aq (45 mL) and extracted with AcOEt (90 mL). The organic layer was washed with brine (30 mL) and dried over $Na_2SO_4$. The solvent was removed under reduced pressure, and the residue was purified by flash chromatography on silica gel ($CHCl_3$ : MeOH : AcOH = 95 : 4 : 1) to yield **12** (84.6 mg, 158 μmol, 26%) as a red solid.

$^1$H NMR (400 MHz; DMSO-d6): δ 11.03 (s, 1H), 8.41 (s, 1H), 8.30 (d, 1H, $J$ = 8.0 Hz), 7.51 (d, 1H, $J$ = 8.0 Hz), 6.77 (d, 1H, $J$ = 8.4 Hz), 6.68 (d, 1H, $J$ = 8.8 Hz).

*Compound **13***

To a stirred solution of **12** (25 mg, 47 μmol) in dry DMF (1.0 mL) was added EDC (14 mg, 73 μmol) and NHS (8.0 mg, 70 μmol). The mixture was stirred at room temperature for 14 h and purified by RP-HPLC (0.1% TFA-AcCN : 0.1% TFA-$H_2O$ = 40 : 60 (0 min), 40 : 60 (5 min), 60 : 40 (25 min)) with a YMC-Pack ODS-A column (5 μm, 250 × 20 mm) at a flow rate of 9.9 mL/min. The target fraction was lyophilised to yield **13** (18 mg, 28 μmol, 61%) as an orange solid.

$^1$H NMR (400 MHz; $CD_3CN/CD_3OD$): δ 8.73 (s, 1H), 8.48 (d, 1H, $J$ = 8.0 Hz), 7.52 (d, 1H, $J$ = 8.0 Hz), 6.74–6.71 (m, 4H, $J$ = 8.4 Hz), 2.94 (s, 4H).

*Compound **4***

To a stirred solution of **6**[S4] (7.0 mg, 8.7 μmol) in dichloromethane (1.0 mL) was added TFA (500 μL). The mixture was stirred at room temperature for 1 h. TFA was removed by co-evaporation with toluene (500 μL) twice. To a stirred solution of this residue in dry DMF (500 μL) was added **13** (8.2 mg, 13 μmol) and DIEA (9.0 μL, 52 μmol). The mixture was stirred at room temperature for 2 h and purified by RP-HPLC (10 mM TEAA-AcCN : 10 mM TEAA-$H_2O$ = 15 : 85 (0 min), 15 : 85 (5 min), 32 : 68 (25 min)) with a YMC-Pack ODS-A column (5 μm, 250 × 20 mm) at a flow rate of 9.9 mL/min. The target fraction was lyophilised to yield **4**

(8 mg, 7.4 μmol, 66%) as an orange oil.

$^1$H NMR (600 MHz; $CD_3OD$): δ 8.47 (s, 1H), 8.18 (s, 1H), 8.00 (d, 1H, $J$ = 7.8 Hz), 7.55 (s, 1H), 7.36–7.33 (m, 2H), 7.11 (s, 1H), 6.93 (d, 2H, $J$ = 9.0 Hz), 6.79–6.75 (m, 2H), 6.63 (d, 2H, $J$ = 9.6 Hz), 6.18 (s, 1H), 4.72 (brs, 2H), 4.57 (t, 2H, $J$ = 4.8 Hz), 4.23 (s, 2H), 3.87 (t, 2H, $J$ = 4.8 Hz), 3.75 (t, 2H, $J$ = 5.4 Hz), 3.63 (t, 2H, $J$ = 5.4 Hz), 3.37 (m, 2H), 2.66 (m, 2H), 2.48–2.44 (m, 4H).

$^{13}$C NMR (150 MHz; $CD_3OD$): δ 177.78, 174.68, 174.65, 174.20, 172.16, 169.25, 157.42, 155.89, 155.38, 149.85, 142.02, 138.50, 136.88, 135.97, 131.64, 130.52, 130.00, 129.22, 126.50, 125.29, 125.11, 123.49, 123.40, 122.90, 122.46, 122.34, 122.25, 117.73, 115.39, 113.38, 110.46, 101.10, 70.56, 69.49, 68.51, 40.99, 39.70, 37.22, 32.50, 32.43, 28.74

HRMS (ESI): Calcd for $(C_{51}H_{42}Br_2F_3N_8O_{15})^+$ $[M+H]^+$: 1221.1083, found: 1221.1053.
